# Taxonomic, ecological and morphological diversity of Ponto-Caspian gammaridean amphipods: a review

**DOI:** 10.1101/2021.01.21.427559

**Authors:** Denis Copilaș-Ciocianu, Dmitry Sidorov

## Abstract

Thanks to its dynamic geological history the Ponto-Caspian region harbors a unique and unusually adaptable fauna, notorious for its invasive species. Gammarid amphipods attained considerable diversity, becoming the world’s second most speciose ancient-lake amphipod radiation. Nonetheless, apart from a few invasive species, this group remains poorly studied. Herein, we review and quantify the taxonomic, morphological and ecological diversity, as well as phylogenetic context of Ponto-Caspian gammarids within the adaptive radiation framework. Published molecular phylogenies indicate that this radiation has a monophyletic mid-Miocene Paratethyan origin, and is nested within the morphologically-conserved Atlanto-Mediterranean genus *Echinogammarus*. We find extensive disparity in body shape, size, ornamentation and appendage length, along a broad ecological gradient from mountain springs to depths exceeding 500 m, on virtually all substrate types (including symbiosis). We propose four putative ecomorphs that appear convergent with distantly related oceanic and Baikal Lake taxa. Thus, the identified patterns support the adaptive radiation model, although extensive further research is needed. A checklist and provisional key to all known endemic species are provided to facilitate taxonomic research. Ponto-Caspian gammarids could be a potentially powerful model for studying adaptive radiations and invasive species evolution.

## Introduction

Ancient lakes are evolutionary cradles, harboring a rich endemic fauna that fascinated biologists for centuries (M. E. Cristescu et al. 2010; Martens 1997). Their confined nature coupled with large size and relative stability over geological time scales promoted lineage accumulation, diversification and ecological specialization. Many of these lineages probably arose through adaptive radiation, an evolutionary process wherein species rapidly evolve from a common ancestor and diversify to occupy various ecological niches (Schluter 2000). Classical examples of adaptive radiations in ancient lakes are cichlid species flocks in African Rift Valley lakes (Salzburger et al. 2014), or the gammarid amphipods inhabiting Lake Baikal (Naumenko et al. 2017).

Situated in the Ponto-Caspian region, the Caspian Sea is the world’s largest ancient lake (M. E. Cristescu et al. 2010). The Azov, Aral and Black seas are also part of this system. These water bodies are remnants of the once widespread epicontinental Paratethys Sea, which stretched from the foothills of the Alps to the Himalayas (Popov et al. 2004). The Paratethys had a turbulent geological history with numerous regression-transgression phases causing drastic salinity fluctuations and repeated episodes of isolation and reconnection with the world ocean (Audzijonyte et al. 2015; Palcu et al. 2019; Popov et al. 2004; Rögl 1999). The uplift of the Caucasus range during the late Miocene triggered the formation and separation of the Black and Caspian seas. During the last two million years these two basins experienced recurrent phases of mutual isolation and reconnection (Krijgsman et al. 2019).

It is thought that this tumultuous geological past drove the evolution of the unusually euryhaline fauna that inhabits the region today (Reid and Orlova 2002). This plasticity has enabled many Ponto-Caspian species to spread across the Northern Hemisphere and become invasive due to human interference (Adrian-Kalchhauser et al. 2020; Cuthbert et al. 2020; Vanderploeg et al. 2002). Nevertheless, many Ponto-Caspian endemics face severe conservation challenges due to climate change, invasive species and multifarious anthropogenic disturbances (Dumont 1995; Gogaladze et al. 2020; Lattuada et al. 2019; Prange et al. 2020). The Ponto-Caspian region is a hot-spot of endemicity and biodiversity with hundreds of species from various animal phyla, but particularly rich in crustaceans (Birstein et al. 1968; Chertoprud et al. 2018; M. E. A. Cristescu and Hebert 2005; Mordukhai-Boltovskoi 1979; Naseka and Bogutskaya 2009; Wesselingh et al. 2019).

Amphipod crustaceans radiated multiple times in the world’s temperate ancient lakes. Several radiations occurred in Lake Titicaca (Hyalellidae) (Adamowicz et al. 2018; Jurado-Rivera et al. 2020), two in Lake Baikal (Gammaridae) (Macdonald et al. 2005; Naumenko et al. 2017), probably two in the Ponto-Caspian basin (Gammaroidea, Corophiidae) (M. E. A. Cristescu and Hebert 2005; Hou et al. 2014), and apparently one radiation in other lakes such as Ohrid (Gammaridae) (Wysocka et al. 2013, 2014), and Fuxian Hu (Anisogammaridae) (Sket and Fišer 2009). Other lakes throughout Asia also harbor endemic species, although their monophyly has yet to be proven. These are Lake Issyk-Kul in Kyrgyzstan (Gammaridae) (Karaman and Pinkster 1977) and Lake Teletskoye (Gammaridae) in Russia (Martynov 1930). In most of these lakes amphipods display a bewildering diversity in form and ecology, with remarkable convergence in body armature among evolutionary and geographically distant groups (Martens 1997).

The endemic amphipod fauna of the Ponto-Caspian basin is one of the world’s most diverse, second only to Lake Baikal (Barnard and Barnard 1983; Väinölä et al. 2008). Among all endemic Ponto-Caspian organisms, amphipods seem to be the most species-rich and successful group, attaining significant ecological and morphological disparity, akin to an adaptive radiation (Derzhavin 1948; Pjatakova and Tarasov 1996; Sars 1895). However, despite these appealing features for evolutionary and ecological studies, Ponto-Caspian amphipods are obscure and poorly known, even ignored in some relatively recent reviews (Martens and Schön 1999). Most attention has been focused on the invasive species that are spreading throughout European freshwaters (e.g. Cristescu et al., 2004; Grabowski et al., 2007; Arbačiauskas et al., 2013; Rewicz et al., 2015), while the non-invasive ones were largely ignored in the last two decades. The taxonomy of the group is rather chaotic due to old and incomplete species descriptions, which led to fuzzy generic diagnoses and lack of a formal system. Even online databases such as World Amphipoda Database (http://www.marinespecies.org/amphipoda/) are incomplete (Horton et al. 2020). Furthermore, a significant part of the literature predates the digital era and is published in Russian, thus not readily available for the international community. As such, to date, there is no comprehensive overview of the Ponto-Caspian amphipod diversity in terms of taxa, ecology and morphology. Some attempts have been made in the past, but these either focused on taxonomy or ecology and never considered the amphipods from all of the Ponto-Caspian basins (Birstein and Romanova 1968; Mordukhai-Boltovskoi 1964, 1979; Pjatakova and Tarasov 1996).

In this study we aim to provide a first comprehensive overview of endemic Ponto-Caspian gammaridean amphipods (taxonomy, morphology and ecology) by examining all of the original species descriptions and relevant literature. Furthermore, by integrating the results of this study with previous phylogenetic research, we strived to identify to which extent the current knowledge on Ponto-Caspian amphipods satisfies the adaptive radiation model (Schluter 2000; Simões et al. 2016). Specifically, we looked for evidence pointing to: I) monophyly of endemic Ponto-Caspian gammarids, an increase in their diversification rates, and III) ecomorphological divergence.

This overview is intended to serve as a foundation and to encourage future eco-evolutionary and taxonomic studies on Ponto-Caspian amphipods. To this end, we also provide a complete checklist and a provisional key to all known endemic species in the hopes of reviving taxonomic interest and to stabilize the systematics of the group.

### Taxonomic diversity

Our study focuses on the Ponto-Caspian amphipod taxa that belong to the superfamily Gammaroidea. Specifically, we included the endemic genera of the family Gammaridae, as well as the fully endemic families Behningiellidae, Caspicolidae, Iphigenellidae, and Pontogammaridae. These taxa form the bulk of the endemic diversity and are most likely a monophyletic group (Hou et al. 2014; Sket and Hou 2018), which is a necessary prerequisite for the adaptive radiation model (Schluter 2000). The remaining Ponto-Caspian endemic amphipods such as *Chelicorophium* (9 spp., Corophiidae), *Gammaracanthus* (1 sp., Gammaracanthidae), *Niphargus* (1 sp.), *Onisimus* (2 spp., Uristidae), and *Monoporeia* (1 sp.) were excluded since they are unrelated to the focal gammarids (Copilaş-Ciocianu et al. 2020; Lowry and Myers 2017; Väinölä et al. 2001). However, we include the monotypic family Caspicolidae because it is very likely a highly derived gammarid (Derzhavin 1944). Although this family is currently included in the infraorder Talitridira by Lowry & Myers (2013), we consider this placement erroneous due to a character coding mistake (see Discussion for further details).

We compiled a checklist of all known Ponto-Caspian gammaroids by reviewing all of the original species descriptions, including re-descriptions. It is presented in Table 1 along with species systematics, native distribution and short taxonomic remarks where necessary. A total of 82 valid extant species are known, belonging to 34 genera and five families: Behningiellidae (3 genera, 4 spp.), Caspicolidae (monotypic), Gammaridae (18 genera, 39 spp.), Iphigenellidae (monotypic) and Pontogammaridae (11 genera, 37 spp.) (Fig. 1a). Five species are doubtful since they may be junior synonyms and further study is needed (Table 1). The most diverse genus is *Pontogammarus* (8 spp.), followed by *Dikerogammarus* and *Obesogammarus* (7 spp. each), *Stenogammarus* (6 spp.), *Chaetogammarus* and *Amathillina* (5 spp. each). Eighteen genera (53%) are monotypic (Fig. 1a). The extinct fossil genera *Andrussovia* (3 spp.) and *Praegmelina* (2 spp.) are currently placed in the Pontogammaridae (Table 1).

**Table 1.**
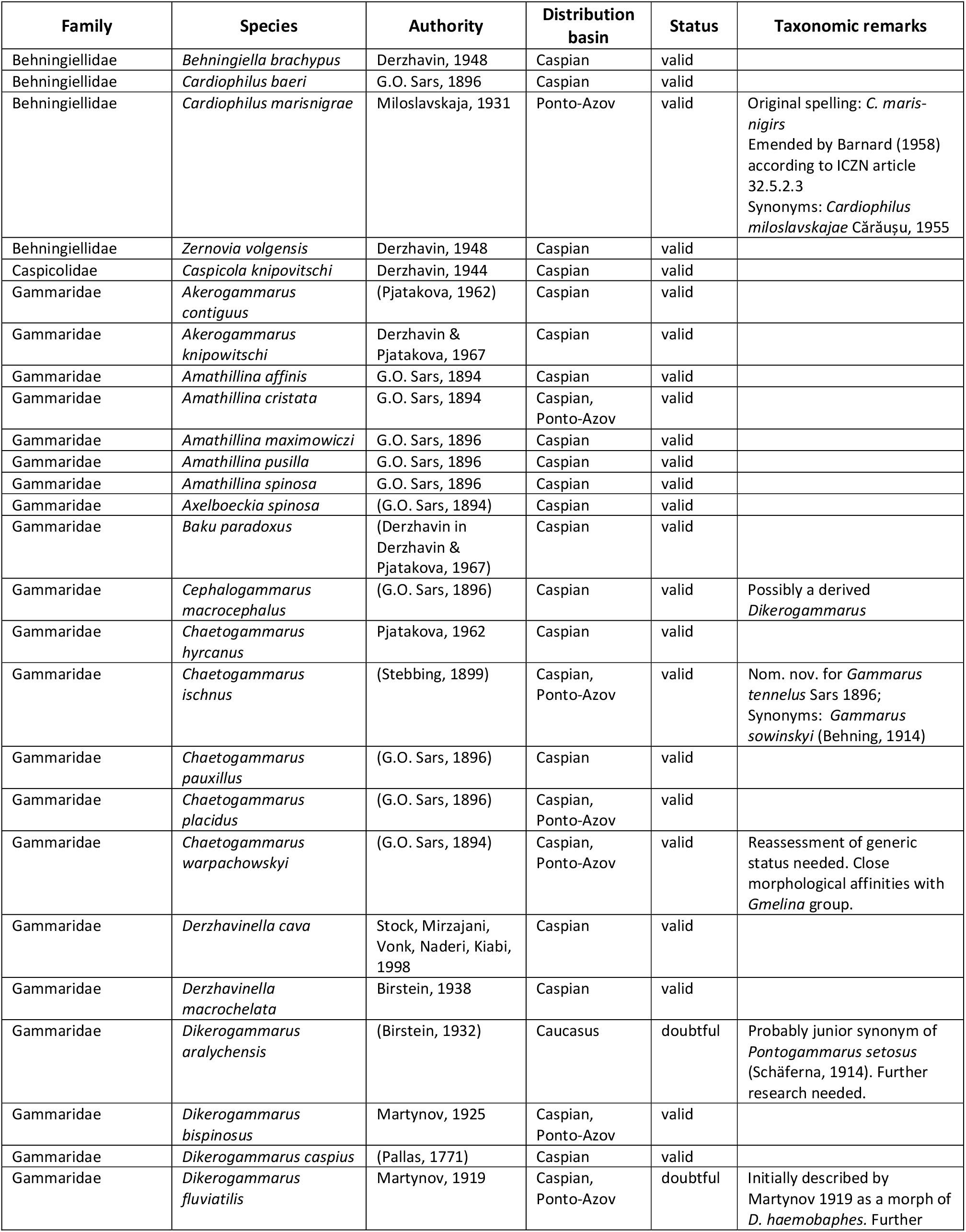
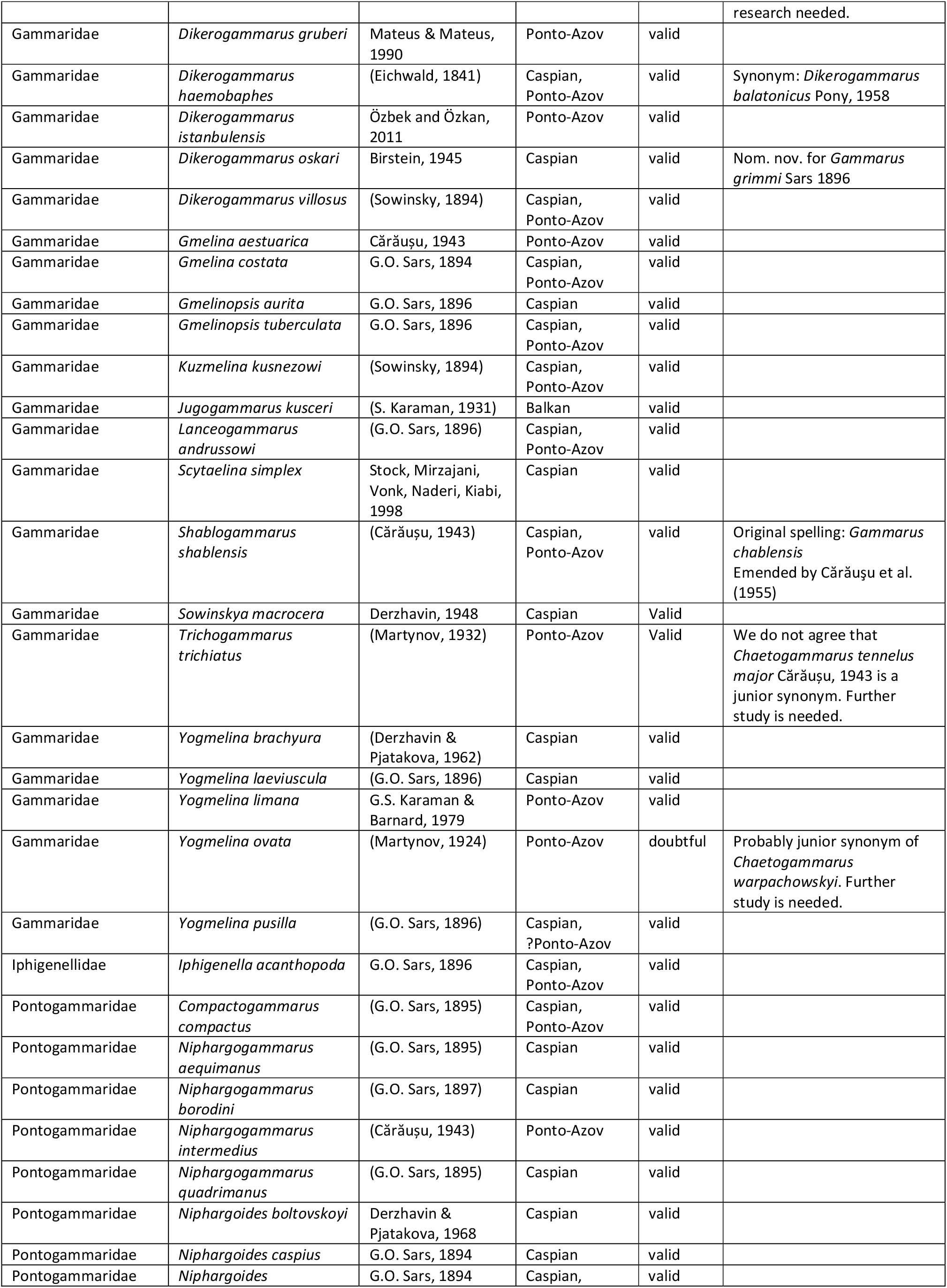
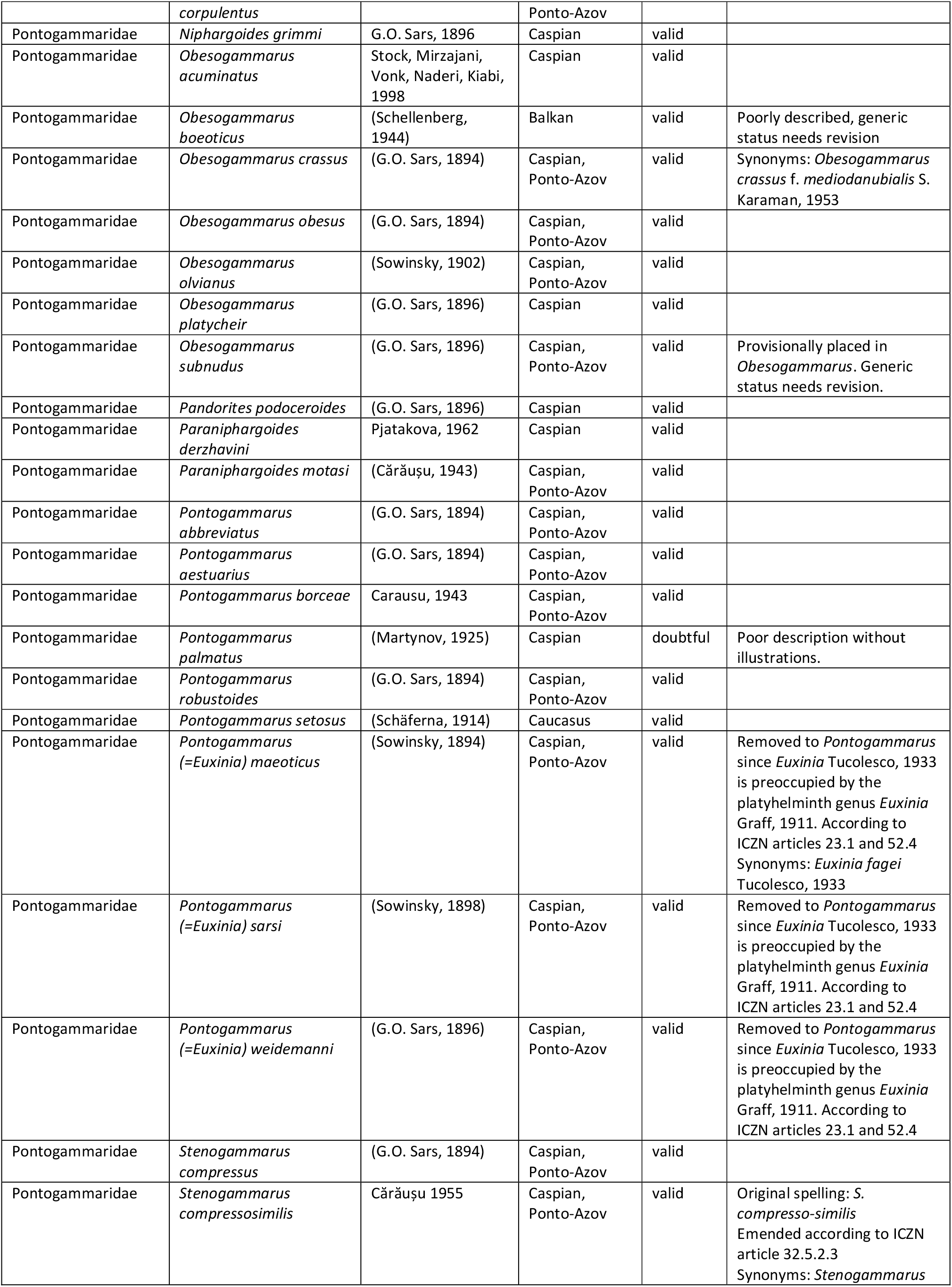
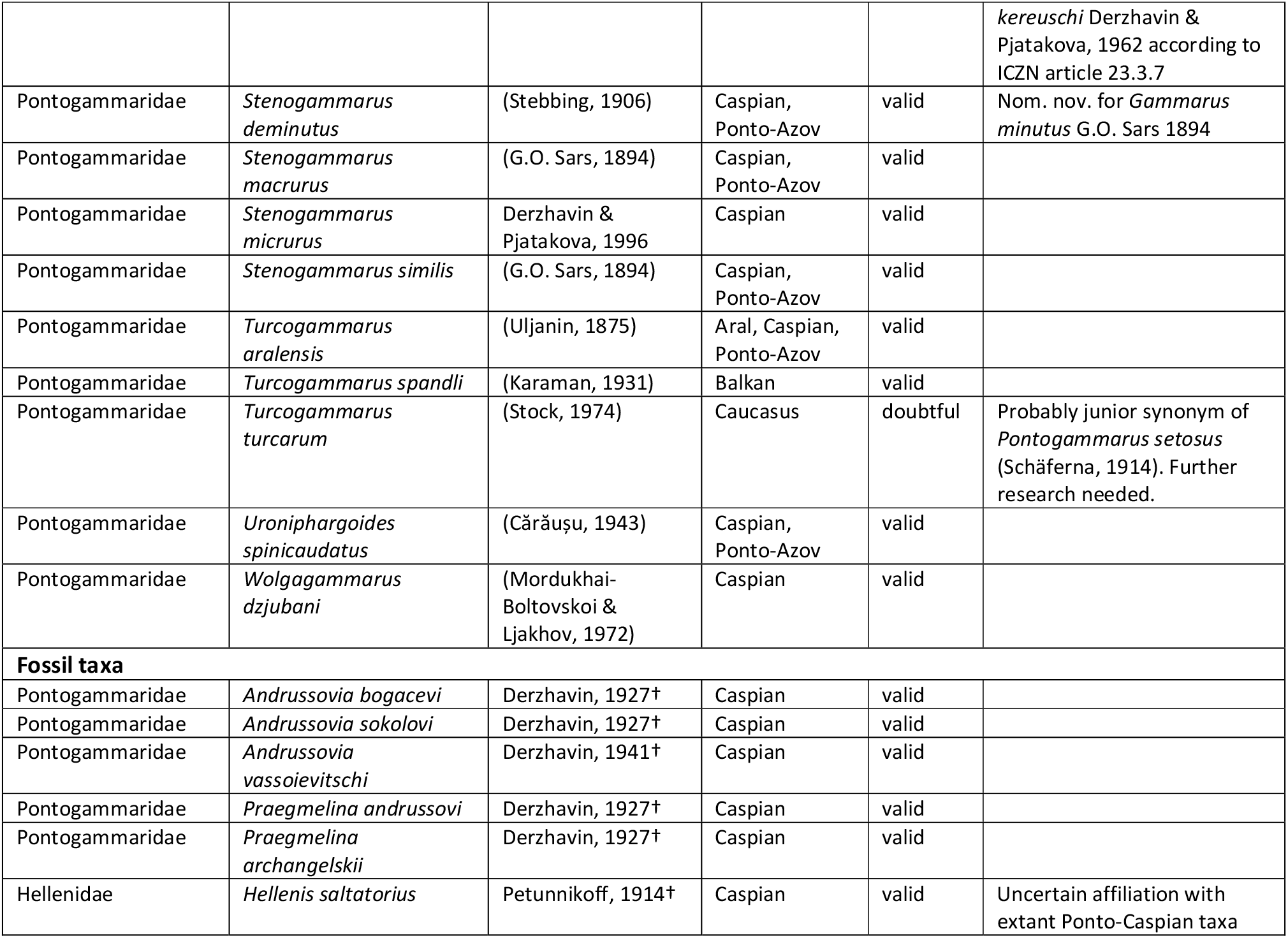
Checklist, taxonomy and native distribution of extant and fossil Ponto-Caspian gammaroid amphipods.

| Family | Species | Authority | Distribution basin | Status | Taxonomic remarks |
| --- | --- | --- | --- | --- | --- |
| Behningiellidae | *Behningiella brachypus* | Derzhavin, 1948 | Caspian | valid |  |
| Behningiellidae | *Cardiophilus baeri* | G.O. Sars, 1896 | Caspian | valid |  |
| Behningiellidae | *Cardiophilus marisnigrae* | Miloslavskaja, 1931 | Ponto-Azov | valid | Original spelling: *C. maris-nigirs*<br>Emended by Barnard (1958) according to ICZN article 32.5.2.3<br>Synonyms: *Cardiophilus miloslavskajae* Cărauşu, 1955 |
| Behningiellidae | *Zernovia volgensis* | Derzhavin, 1948 | Caspian | valid |  |
| Caspicolidae | *Caspicola knipovitschi* | Derzhavin, 1944 | Caspian | valid |  |
| Gammaridae | *Akerogammarus contiguus* | (Pjatakova, 1962) | Caspian | valid |  |
| Gammaridae | *Akerogammarus knipowitschi* | Derzhavin & Pjatakova, 1967 | Caspian | valid |  |
| Gammaridae | *Amathillina affinis* | G.O. Sars, 1894 | Caspian | valid |  |
| Gammaridae | *Amathillina cristata* | G.O. Sars, 1894 | Caspian, Ponto-Azov | valid |  |
| Gammaridae | *Amathillina maximowiczi* | G.O. Sars, 1896 | Caspian | valid |  |
| Gammaridae | *Amathillina pusilla* | G.O. Sars, 1896 | Caspian | valid |  |
| Gammaridae | *Amathillina spinosa* | G.O. Sars, 1896 | Caspian | valid |  |
| Gammaridae | *Axelboeckia spinosa* | (G.O. Sars, 1894) | Caspian | valid |  |
| Gammaridae | *Baku paradoxus* | (Derzhavin in Derzhavin & Pjatakova, 1967) | Caspian | valid |  |
| Gammaridae | *Cephalogammarus macrocephalus* | (G.O. Sars, 1896) | Caspian | valid | Possibly a derived *Dikerogammarus* |
| Gammaridae | *Chaetogammarus hyrcanus* | Pjatakova, 1962 | Caspian | valid |  |
| Gammaridae | *Chaetogammarus ischnus* | (Stebbing, 1899) | Caspian, Ponto-Azov | valid | Nom. nov. for *Gammarus tennelus* Sars 1896; Synonyms: *Gammarus sowinskyi* (Behning, 1914) |
| Gammaridae | *Chaetogammarus pauxillus* | (G.O. Sars, 1896) | Caspian | valid |  |
| Gammaridae | *Chaetogammarus placidus* | (G.O. Sars, 1896) | Caspian, Ponto-Azov | valid |  |
| Gammaridae | *Chaetogammarus warpachowskyi* | (G.O. Sars, 1894) | Caspian, Ponto-Azov | valid | Reassessment of generic status needed. Close morphological affinities with *Gmelina* group. |
| Gammaridae | *Derzhavinella cava* | Stock, Mirzajani, Vonk, Naderi, Kiabi, 1998 | Caspian | valid |  |
| Gammaridae | *Derzhavinella macrochelata* | Birstein, 1938 | Caspian | valid |  |
| Gammaridae | *Dikerogammarus aralychensis* | (Birstein, 1932) | Caucasus | doubtful | Probably junior synonym of *Pontogammarus setosus* (Schäferna, 1914). Further research needed. |
| Gammaridae | *Dikerogammarus bispinosus* | Martynov, 1925 | Caspian, Ponto-Azov | valid |  |
| Gammaridae | *Dikerogammarus caspius* | (Pallas, 1771) | Caspian | valid |  |
| Gammaridae | *Dikerogammarus fluviatilis* | Martynov, 1919 | Caspian, Ponto-Azov | doubtful | Initially described by Martynov 1919 as a morph of *D. haemobaphes*. Further research needed. |
| Gammaridae | *Dikerogammarus gruberi* | Mateus & Mateus, 1990 | Ponto-Azov | valid |  |
| Gammaridae | *Dikerogammarus haemobaphes* | (Eichwald, 1841) | Caspian, Ponto-Azov | valid | Synonym: *Dikerogammarus balatonicus* Pony, 1958 |
| Gammaridae | *Dikerogammarus istanbulensis* | Özbek and Özkan, 2011 | Ponto-Azov | valid |  |
| Gammaridae | *Dikerogammarus oskari* | Birstein, 1945 | Caspian | valid | Nom. nov. for *Gammarus grimmi* Sars 1896 |
| Gammaridae | *Dikerogammarus villosus* | (Sowinsky, 1894) | Caspian, Ponto-Azov | valid |  |
| Gammaridae | *Gmelina aestuarica* | Cărașu, 1943 | Ponto-Azov | valid |  |
| Gammaridae | *Gmelina costata* | G.O. Sars, 1894 | Caspian, Ponto-Azov | valid |  |
| Gammaridae | *Gmelinopsis aurita* | G.O. Sars, 1896 | Caspian | valid |  |
| Gammaridae | *Gmelinopsis tuberculata* | G.O. Sars, 1896 | Caspian, Ponto-Azov | valid |  |
| Gammaridae | *Kuzmelina kusnezowi* | (Sowinsky, 1894) | Caspian, Ponto-Azov | valid |  |
| Gammaridae | *Jugogammarus kusceri* | (S. Karaman, 1931) | Balkan | valid |  |
| Gammaridae | *Lanceogammarus andrussowi* | (G.O. Sars, 1896) | Caspian, Ponto-Azov | valid |  |
| Gammaridae | *Scytaelina simplex* | Stock, Mirzajani, Vonk, Naderi, Kiabi, 1998 | Caspian | valid |  |
| Gammaridae | *Shablogammarus shablensis* | (Cărașu, 1943) | Caspian, Ponto-Azov | valid | Original spelling: *Gammarus chablensis*<br>Emended by Cărașu et al. (1955) |
| Gammaridae | *Sowinskyia macrocera* | Derzhavin, 1948 | Caspian | Valid |  |
| Gammaridae | *Trichogammarus trichiatus* | (Martynov, 1932) | Ponto-Azov | Valid | We do not agree that *Chaetogammarus tennelus major* Cărașu, 1943 is a junior synonym. Further study is needed. |
| Gammaridae | *Yogmelina brachyura* | (Derzhavin & Pjatakova, 1962) | Caspian | valid |  |
| Gammaridae | *Yogmelina laeviuscula* | (G.O. Sars, 1896) | Caspian | valid |  |
| Gammaridae | *Yogmelina limana* | G.S. Karaman & Barnard, 1979 | Ponto-Azov | valid |  |
| Gammaridae | *Yogmelina ovata* | (Martynov, 1924) | Ponto-Azov | doubtful | Probably junior synonym of *Chaetogammarus warpachowskyi*. Further study is needed. |
| Gammaridae | *Yogmelina pusilla* | (G.O. Sars, 1896) | Caspian, ?Ponto-Azov | valid |  |
| Iphigenellidae | *Iphigenella acanthopoda* | G.O. Sars, 1896 | Caspian, Ponto-Azov | valid |  |
| Pontogammaridae | *Compactogammarus compactus* | (G.O. Sars, 1895) | Caspian, Ponto-Azov | valid |  |
| Pontogammaridae | *Niphargogammarus aequimanus* | (G.O. Sars, 1895) | Caspian | valid |  |
| Pontogammaridae | *Niphargogammarus borodini* | (G.O. Sars, 1897) | Caspian | valid |  |
| Pontogammaridae | *Niphargogammarus intermedius* | (Cărașu, 1943) | Ponto-Azov | valid |  |
| Pontogammaridae | *Niphargogammarus quadrimanus* | (G.O. Sars, 1895) | Caspian | valid |  |
| Pontogammaridae | *Niphargoides boltovskoyi* | Derzhavin & Pjatakova, 1968 | Caspian | valid |  |
| Pontogammaridae | *Niphargoides caspius* | G.O. Sars, 1894 | Caspian | valid |  |
| Pontogammaridae | *Niphargoides* | G.O. Sars, 1894 | Caspian, | valid |  |
|  | *corpulentus* |  | Ponto-Azov |  |  |
| Pontogammaridae | *Niphargoides grimmi* | G.O. Sars, 1896 | Caspian | valid |  |
| Pontogammaridae | *Obesogammarus acuminatus* | Stock, Mirzajani, Vonk, Naderi, Kiabi, 1998 | Caspian | valid |  |
| Pontogammaridae | *Obesogammarus boeoticus* | (Schellenberg, 1944) | Balkan | valid | Poorly described, generic status needs revision |
| Pontogammaridae | *Obesogammarus crassus* | (G.O. Sars, 1894) | Caspian, Ponto-Azov | valid | Synonyms: *Obesogammarus crassus* f. *mediodanubialis* S. Karaman, 1953 |
| Pontogammaridae | *Obesogammarus obesus* | (G.O. Sars, 1894) | Caspian, Ponto-Azov | valid |  |
| Pontogammaridae | *Obesogammarus olivianus* | (Sowinsky, 1902) | Caspian, Ponto-Azov | valid |  |
| Pontogammaridae | *Obesogammarus platycheir* | (G.O. Sars, 1896) | Caspian | valid |  |
| Pontogammaridae | *Obesogammarus subnudus* | (G.O. Sars, 1896) | Caspian, Ponto-Azov | valid | Provisionally placed in *Obesogammarus*. Generic status needs revision. |
| Pontogammaridae | *Pandorites podoceroideis* | (G.O. Sars, 1896) | Caspian | valid |  |
| Pontogammaridae | *Paraniphargoides derzhavini* | Pjatakova, 1962 | Caspian | valid |  |
| Pontogammaridae | *Paraniphargoides motasi* | (Cărbăușu, 1943) | Caspian, Ponto-Azov | valid |  |
| Pontogammaridae | *Pontogammarus abbreviatus* | (G.O. Sars, 1894) | Caspian, Ponto-Azov | valid |  |
| Pontogammaridae | *Pontogammarus aestuarius* | (G.O. Sars, 1894) | Caspian, Ponto-Azov | valid |  |
| Pontogammaridae | *Pontogammarus boraceae* | Caraus, 1943 | Caspian, Ponto-Azov | valid |  |
| Pontogammaridae | *Pontogammarus palmatus* | (Martynov, 1925) | Caspian | doubtful | Poor description without illustrations. |
| Pontogammaridae | *Pontogammarus robustoides* | (G.O. Sars, 1894) | Caspian, Ponto-Azov | valid |  |
| Pontogammaridae | *Pontogammarus setosus* | (Schäferna, 1914) | Caucasus | valid |  |
| Pontogammaridae | *Pontogammarus (=Euxinia) maeoticus* | (Sowinsky, 1894) | Caspian, Ponto-Azov | valid | Removed to *Pontogammarus* since *Euxinia* Tucolesco, 1933 is preoccupied by the platyhelminth genus *Euxinia* Graff, 1911. According to ICZN articles 23.1 and 52.4 Synonyms: *Euxinia fagei* Tucolesco, 1933 |
| Pontogammaridae | *Pontogammarus (=Euxinia) sarsi* | (Sowinsky, 1898) | Caspian, Ponto-Azov | valid | Removed to *Pontogammarus* since *Euxinia* Tucolesco, 1933 is preoccupied by the platyhelminth genus *Euxinia* Graff, 1911. According to ICZN articles 23.1 and 52.4 |
| Pontogammaridae | *Pontogammarus (=Euxinia) weidmanni* | (G.O. Sars, 1896) | Caspian, Ponto-Azov | valid | Removed to *Pontogammarus* since *Euxinia* Tucolesco, 1933 is preoccupied by the platyhelminth genus *Euxinia* Graff, 1911. According to ICZN articles 23.1 and 52.4 |
| Pontogammaridae | *Stenogammarus compressus* | (G.O. Sars, 1894) | Caspian, Ponto-Azov | valid |  |
| Pontogammaridae | *Stenogammarus compressosimilis* | Cărbăușu 1955 | Caspian, Ponto-Azov | valid | Original spelling: *S. compresso-similis* Emended according to ICZN article 32.5.2.3 Synonyms: *Stenogammarus* *kereuschi* Derzhavin & Pjatakova, 1962 according to ICZN article 23.3.7 |
| Pontogammaridae | *Stenogammarus deminutus* | (Stebbing, 1906) | Caspian, Ponto-Azov | valid | Nom. nov. for *Gammarus minutus* G.O. Sars 1894 |
| Pontogammaridae | *Stenogammarus macrurus* | (G.O. Sars, 1894) | Caspian, Ponto-Azov | valid |  |
| Pontogammaridae | *Stenogammarus micrurus* | Derzhavin & Pjatakova, 1996 | Caspian | valid |  |
| Pontogammaridae | *Stenogammarus similis* | (G.O. Sars, 1894) | Caspian, Ponto-Azov | valid |  |
| Pontogammaridae | *Turcogammarus aralensis* | (Uljanin, 1875) | Aral, Caspian, Ponto-Azov | valid |  |
| Pontogammaridae | *Turcogammarus spandli* | (Karaman, 1931) | Balkan | valid |  |
| Pontogammaridae | *Turcogammarus turcarum* | (Stock, 1974) | Caucasus | doubtful | Probably junior synonym of *Pontogammarus setosus* (Schäferna, 1914). Further research needed. |
| Pontogammaridae | *Uroniphargoides spinicaudatus* | (Cărbăușu, 1943) | Caspian, Ponto-Azov | valid |  |
| Pontogammaridae | *Wolgagammarus dzjubani* | (Mordukhai-Boltovskoi & Ljakhov, 1972) | Caspian | valid |  |
| **Fossil taxa** |  |  |  |  |  |
| Pontogammaridae | *Andrussovia bogacevi* | Derzhavin, 1927† | Caspian | valid |  |
| Pontogammaridae | *Andrussovia sokolovi* | Derzhavin, 1927† | Caspian | valid |  |
| Pontogammaridae | *Andrussovia vassoievitschi* | Derzhavin, 1941† | Caspian | valid |  |
| Pontogammaridae | *Praegmelina andrussovi* | Derzhavin, 1927† | Caspian | valid |  |
| Pontogammaridae | *Praegmelina archangelskii* | Derzhavin, 1927† | Caspian | valid |  |
| Hellenidae | *Hellenis saltatorius* | Petunnikoff, 1914† | Caspian | valid | Uncertain affiliation with extant Ponto-Caspian taxa |

**Fig. 1.**
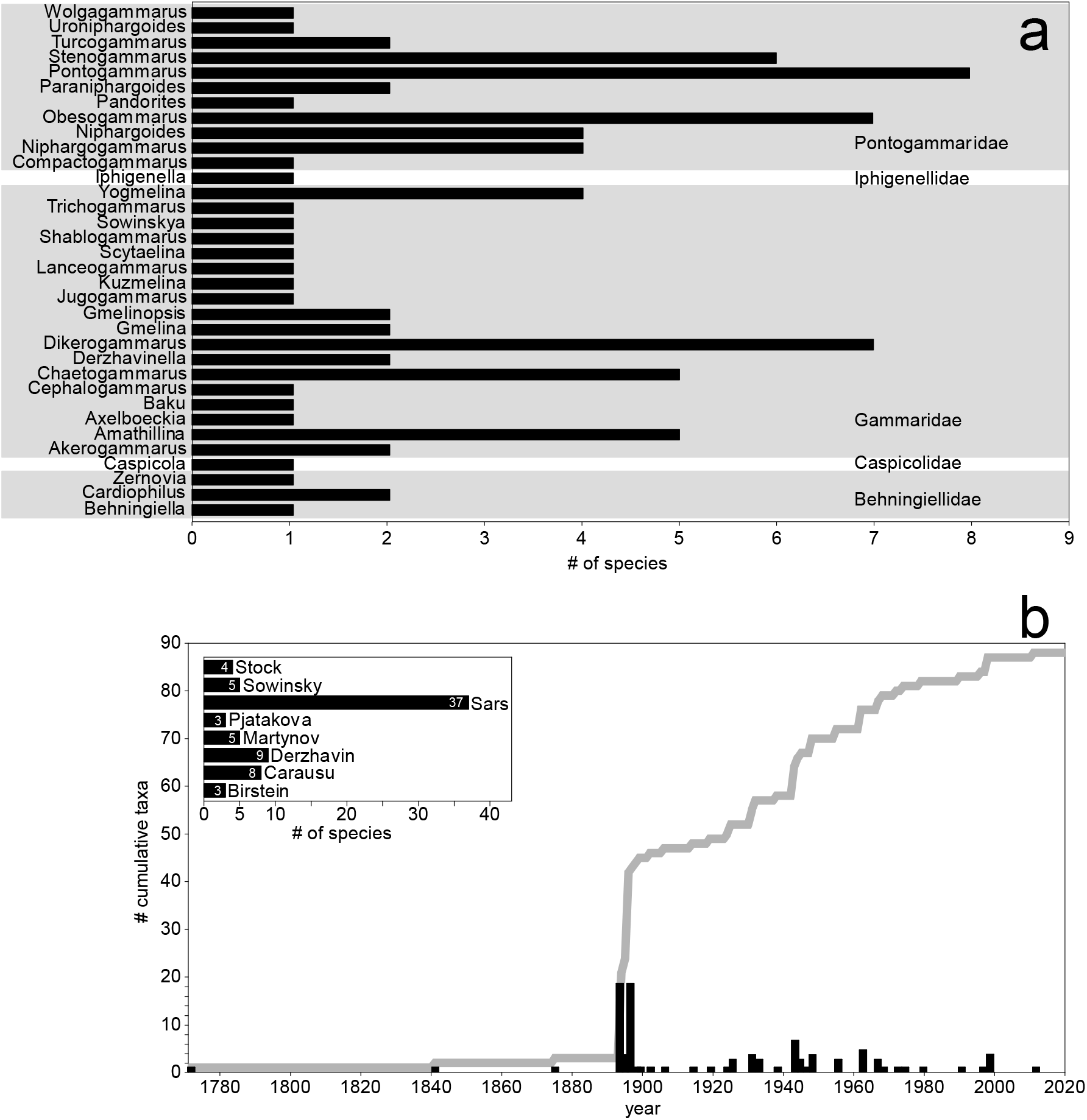
a) Species richness within genera and families. Only valid and extant species were considered. b) Trends in species descriptions through time. The thick gray line indicates the cumulative number of species while black bars indicate the number of species described in that respective year. The inset graph depicts the number of species described by the most prominent authors

The trend of species description through time reveals little taxonomic activity from the 18^th^ to late 19^th^ centuries, a sudden increase with Sars’ monographs in the late 19^th^ century, followed by a more or less steady increase towards the present day with peaks of activity in the middle 20^th^ century by Russian and Romanian authors (Fig. 1b). A noticeable stagnation can be observed in the last two decades. By far the most prolific author was G. O. Sars (37 spp.), followed by A. N. Derzhavin (9 spp.) and S. Cărăuşu (8 spp.)(Fig. 1b inset).

A provisional key to all known endemic families, genera and species (including non-Gammaroidea) is provided in the Appendix. We emphasize that some taxa are poorly known and have an uncertain generic placement.

### Morphology

To explore morphological diversity we extracted data only from those original species descriptions or re-descriptions that provided good quality habitus illustrations (73% of all species) (Cărăuşu 1943; Cărăuşu et al. 1955; Derzhavin 1944, 1948; Sars 1894a, 1894b, 1895, 1896). This was necessary because we used the ratios of various body parts and appendages to total body length. In total, we calculated ratios for 53 traits reflecting as much as possible the overall body shape and functional morphology (see Supplementary information Tables S1-S2 and Fig. S1) (Fišer et al. 2009). The ratios were measured using the Digimizer software (https://www.digimizer.com/). Whenever possible, both sexes were included. We acknowledge that these illustration obtained ratios do not provide the most exhaustive nor precise morphological detail. However, given that these data have a broad taxonomic coverage, we consider this analysis as a crucial preliminary step in quantifying and understanding the morphological diversity of Ponto-Caspian amphipods.

Apart from ratios, we also extracted body size information from the literature and included it in the analysis as well. The 53 ratio + body-size dataset was subjected to a Principal Component Analysis (PCA) based on a correlation matrix to visualize morphological gradients and similarity among genera. Analysis was performed using Statistica 8.0 (StatSoft, Inc.,Tulsa, OK, USA).

We find substantial diversity in body shape and size. The habitus of representative species is presented in Fig. 2. Body size varies by almost an order of magnitude (3.5 to 27 mm) (Figs. 2, 6). The PCA plot indicates significant morphological disparity (Fig. 3). The first four PCA axes explained 22.76, 14.12, 10.17 and 9.28% (56.34%) of the total variation. The first principal component separated species along a gradient from stout bodies with deep coxae and short antennae to slender bodies with shallow coxae and long antennae (Fig. 3a). The second principal component distinguished a gradient along which species were separated by the length of walking appendages and the depth of the tergum (Fig. 3a). The loadings of traits on the PCA axes are presented in Supplementary information Table 3.

**Fig. 2.**
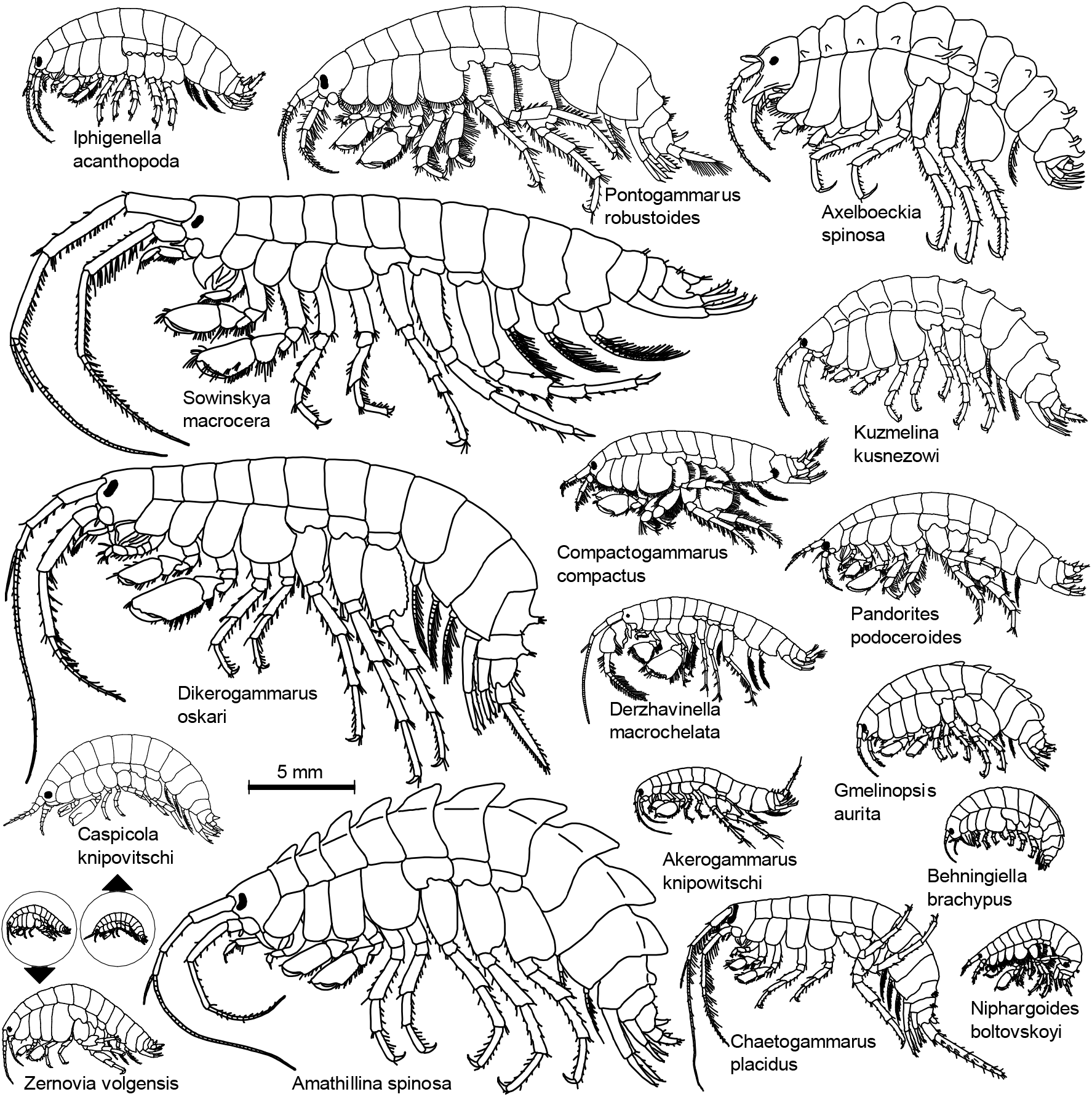
Habitus and morphological diversity of Ponto-Caspian amphipods. *Caspicola knipovitschi* and *Zernovia volgensis* are shown to scale in circles and enlarged outside the circles. All images are redrawn after the original

**Fig. 3.**
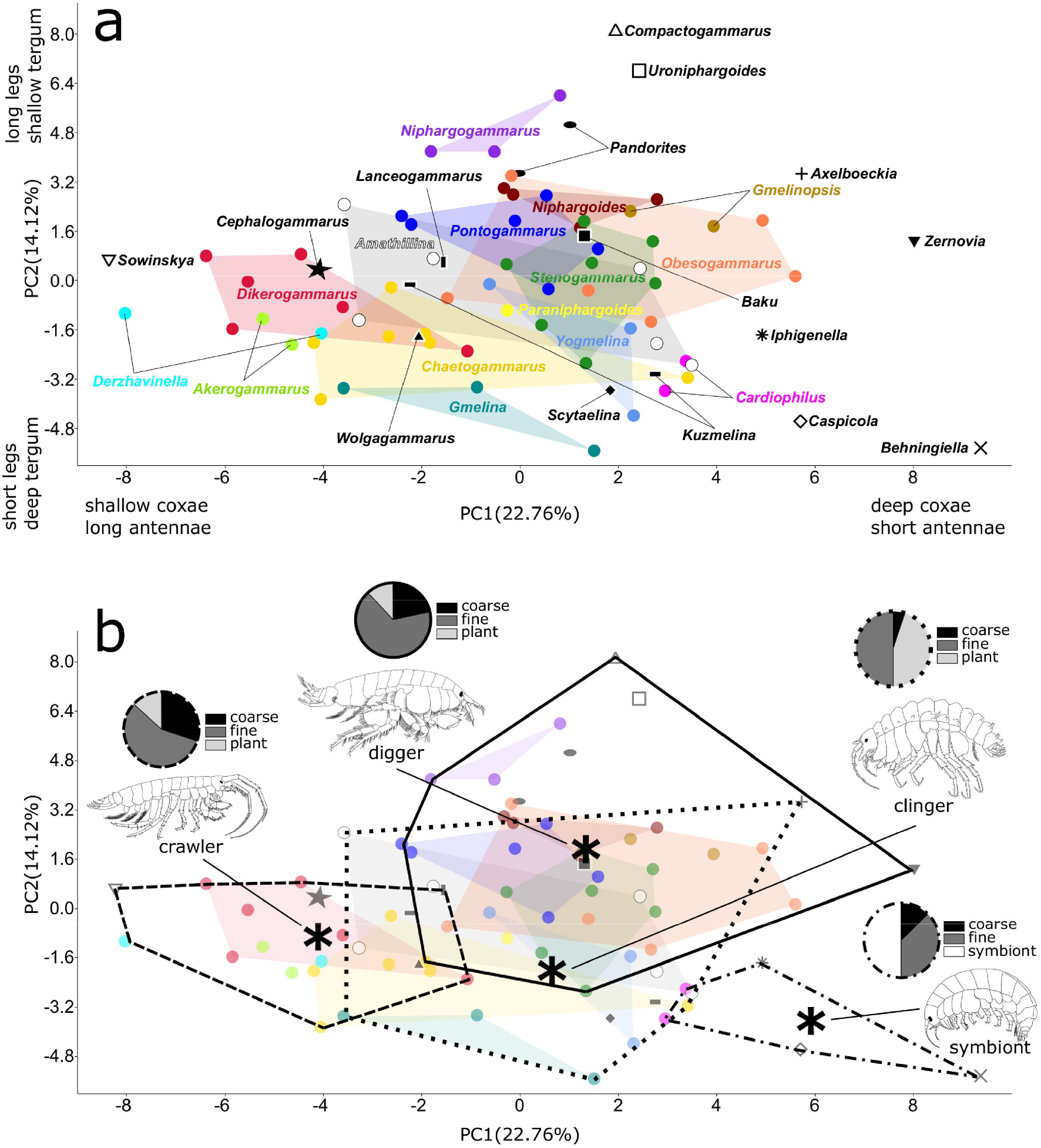
a) PCA scatterplot depicting the morphological gradients along the first two axes. Genera represented by at least three data points are shown with a uniquely colored convex hull and dots. Monotypic genera are depicted with various black symbols and shapes. b) The same PCA as in a) but with convex hulls delineating putative ecomorphs. Asterisks indicate morph centroid. For each morph a representative species is shown. The pie-charts indicate the proportion of species occurring on various substrates within each ecomorph

**Fig. 4.**
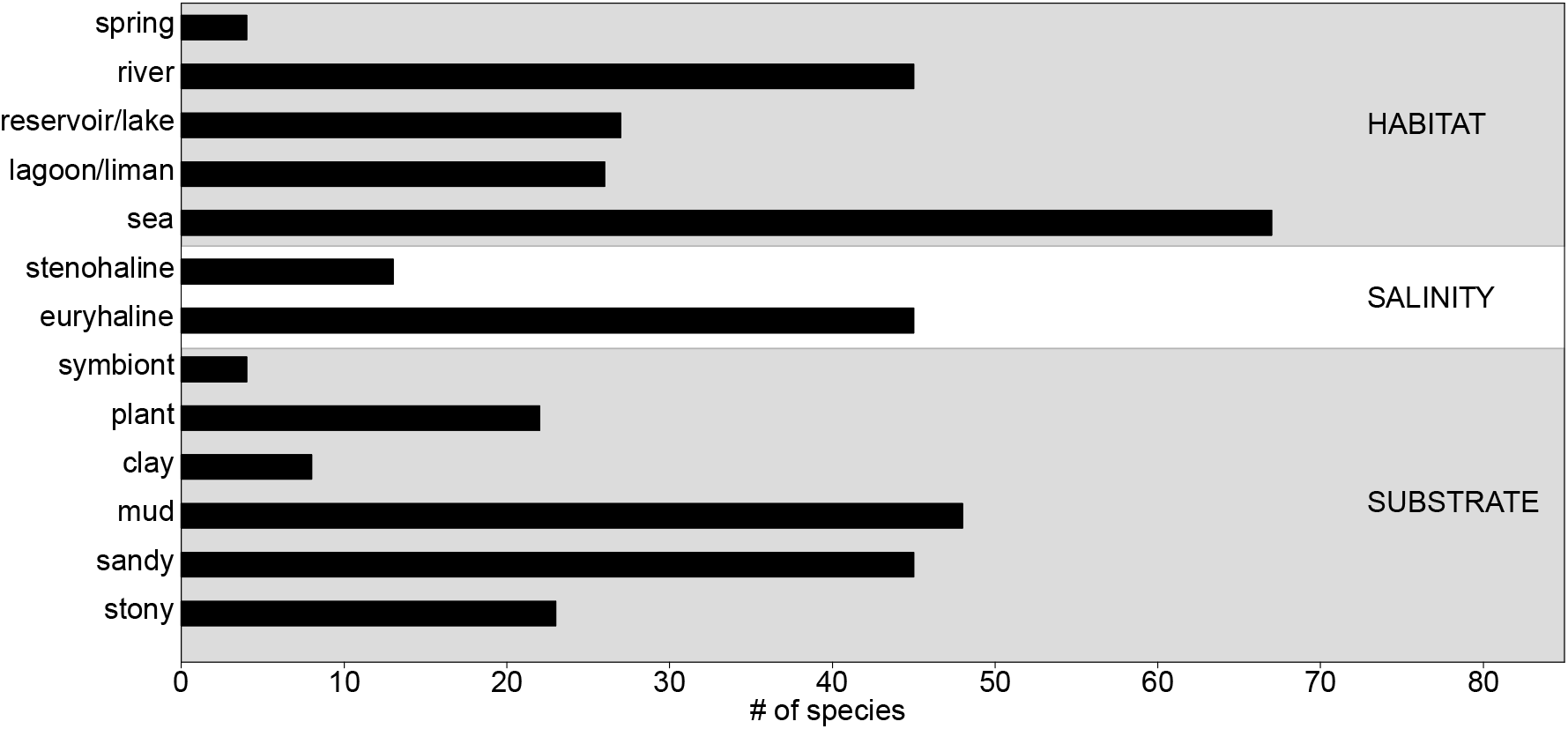
Number of species occurring on various substrates, habitats and salinities

**Fig. 5.**
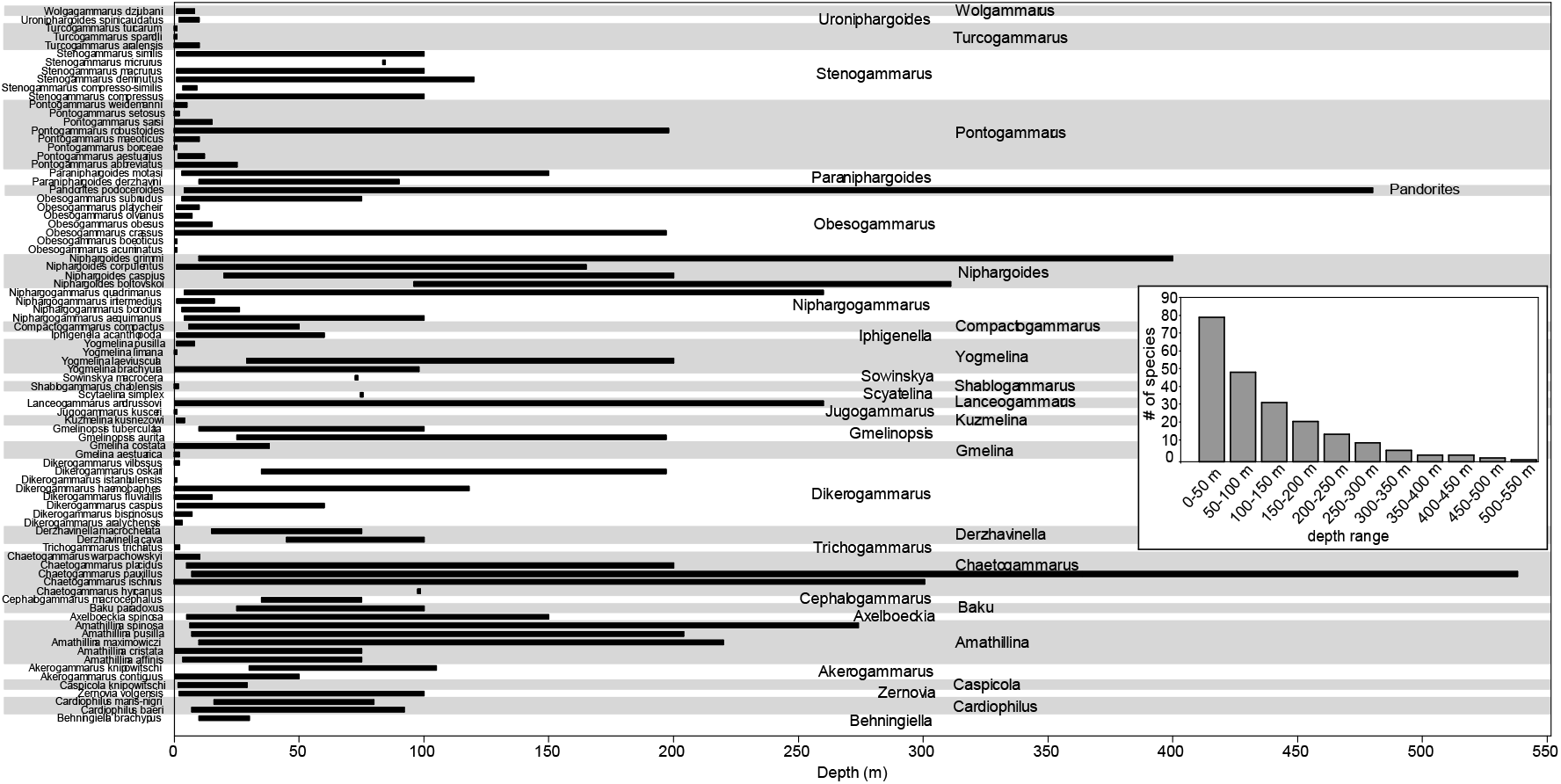
Depth ranges structured by taxonomic composition. The inset graph depicts the number of species occurring in 50 m depth intervals

**Fig. 6.**
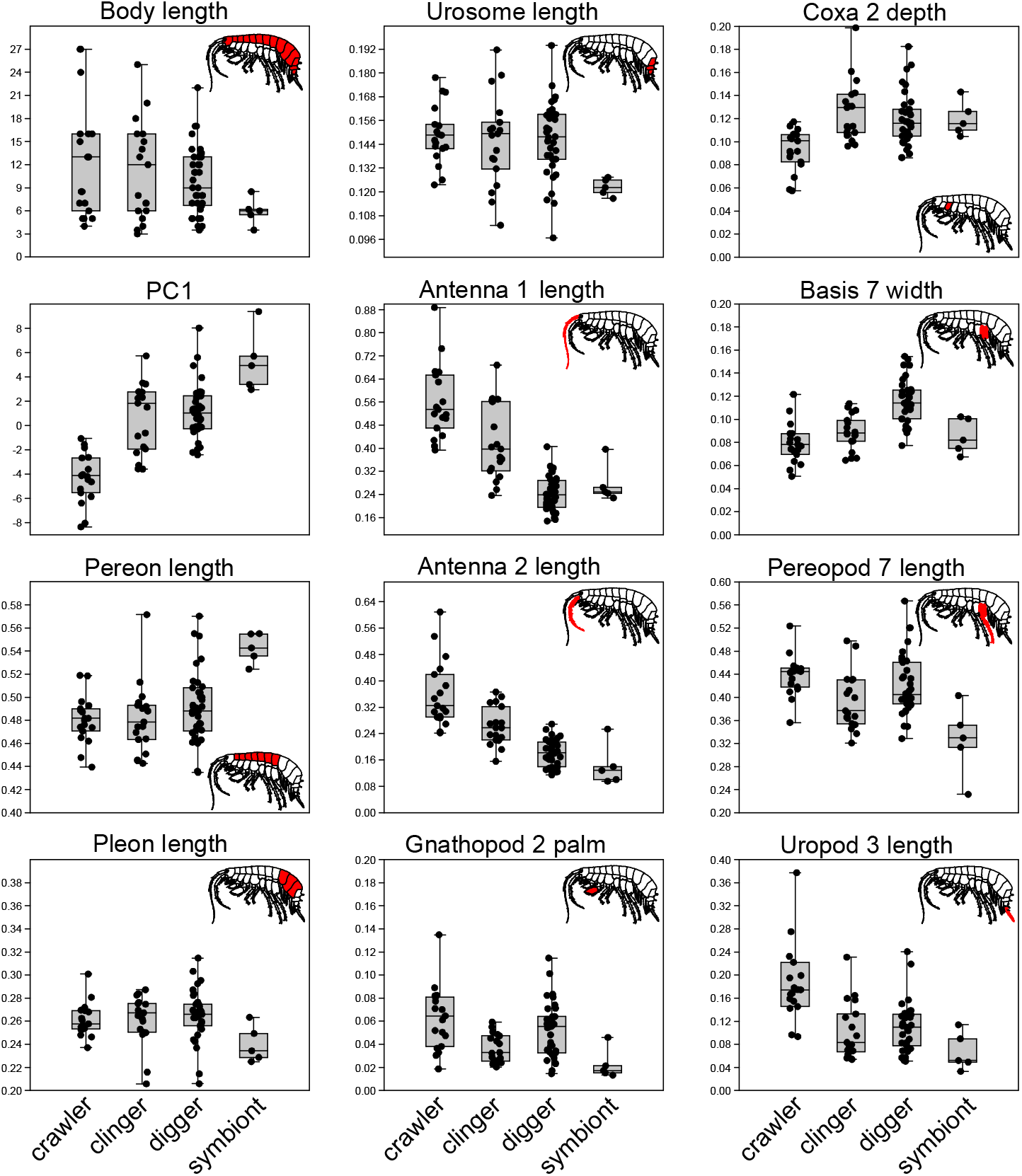
Boxplots comparing selected traits among the four proposed ecomorphs. PC1 refers to the first principal component resulting from the PCA analysis. It mainly describes the gradient from slender bodies with long antennae (negative values) to stout bodies with short antennae (positive values). All traits except body length and PC1 values are presented relative to total body length

There is significant variation with respect to body armature as well. Although most species are generally smooth, there are diverse patterns of ornamentation with either a medial keel that extends throughout different body regions (e.g. *Amathillina, Gmelina* and *Gmelinopsis*) to double dorso-lateral cuspidation (*Kuzmelina*), to lateral spines and dorsal protuberances (*Axelboeckia*) (Fig. 2).

Most genera seem to be relatively well defined in morphospace. However, *Amathillina* and *Obesogammarus* overlap broadly with other genera (Fig. 3a). The monotypic genera (shown with black and white symbols in Fig. 3a) are generally distinct from the more speciose ones, often lying towards the extreme ends of the morphological gradients.

### Ecology

To provide a synopsis of ecology we reviewed all the original species descriptions and the relevant literature (Birstein and Romanova 1968; Pjatakova and Tarasov 1996). We gathered data regarding depth (minimum and maximum), habitat (sea, lagoon, lake/reservoir, river and spring), salinity (steno- and/or euryhaline) and substrate type (stone, sand, mud, clay, plant and symbiotic relationships).

Our review highlights important ecological diversity within the Ponto-Caspian radiation. With respect to habitat, most species live in the sea (67 spp.) and lower courses of rivers (45 spp.), followed by brackish lagoons (26 spp.) and freshwater lakes or reservoirs (27 spp.). Only four species occur exclusively in springs and streams (Table 2, Fig. 4). With respect to salinity, it appears that most species are euryhaline, tolerating both fresh as well as brackish waters. However, salinity preference is not known for many species. With respect to substrate, the great majority of species occur on sandy and muddy substrates, followed by stones and plants. Four species seem to be associated with other organisms such as bivalve mollusks and crayfish (Table 2, Fig. 4). All of the ecological data is summarized in Table 2.

**Table 2.**
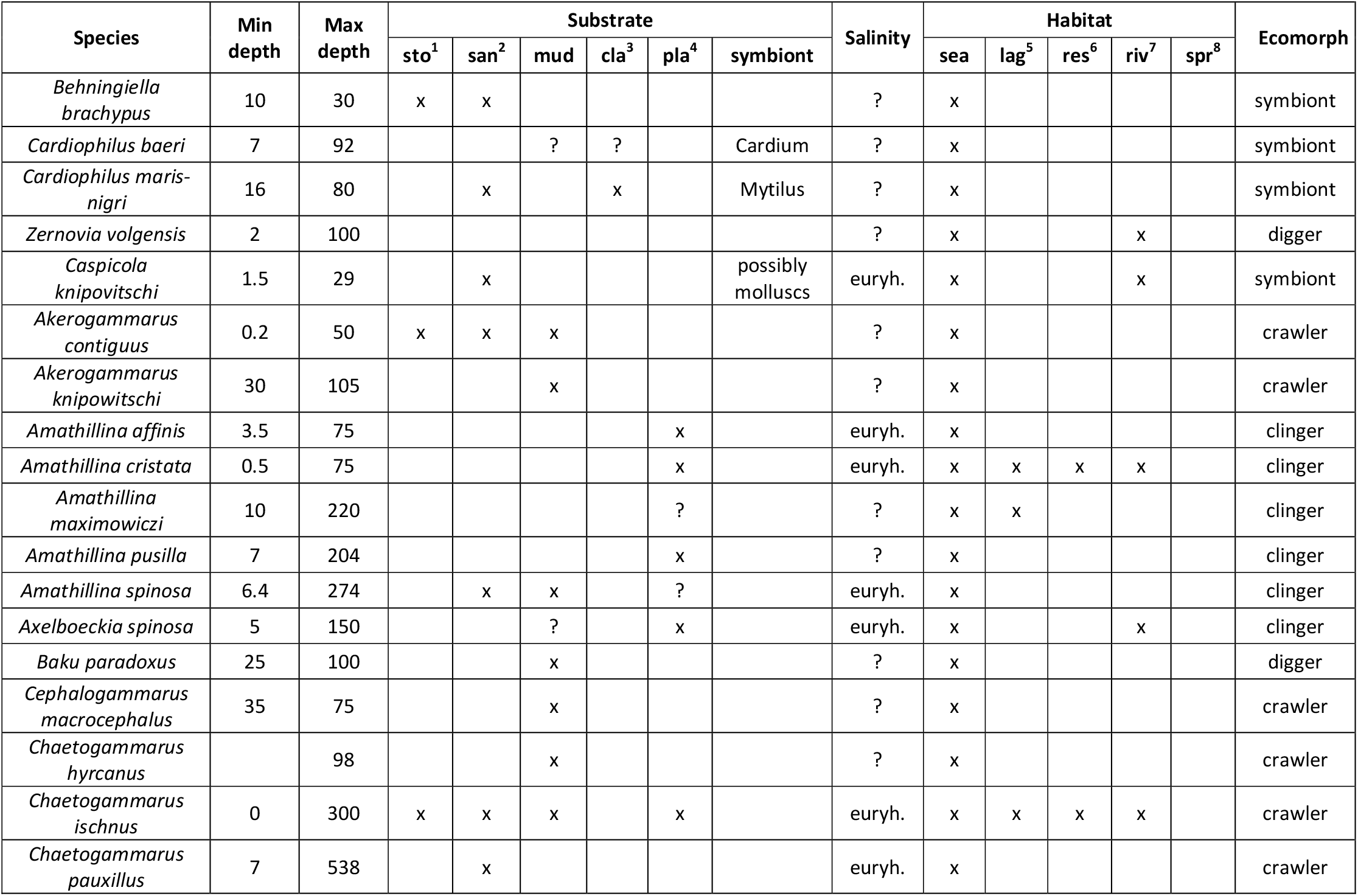
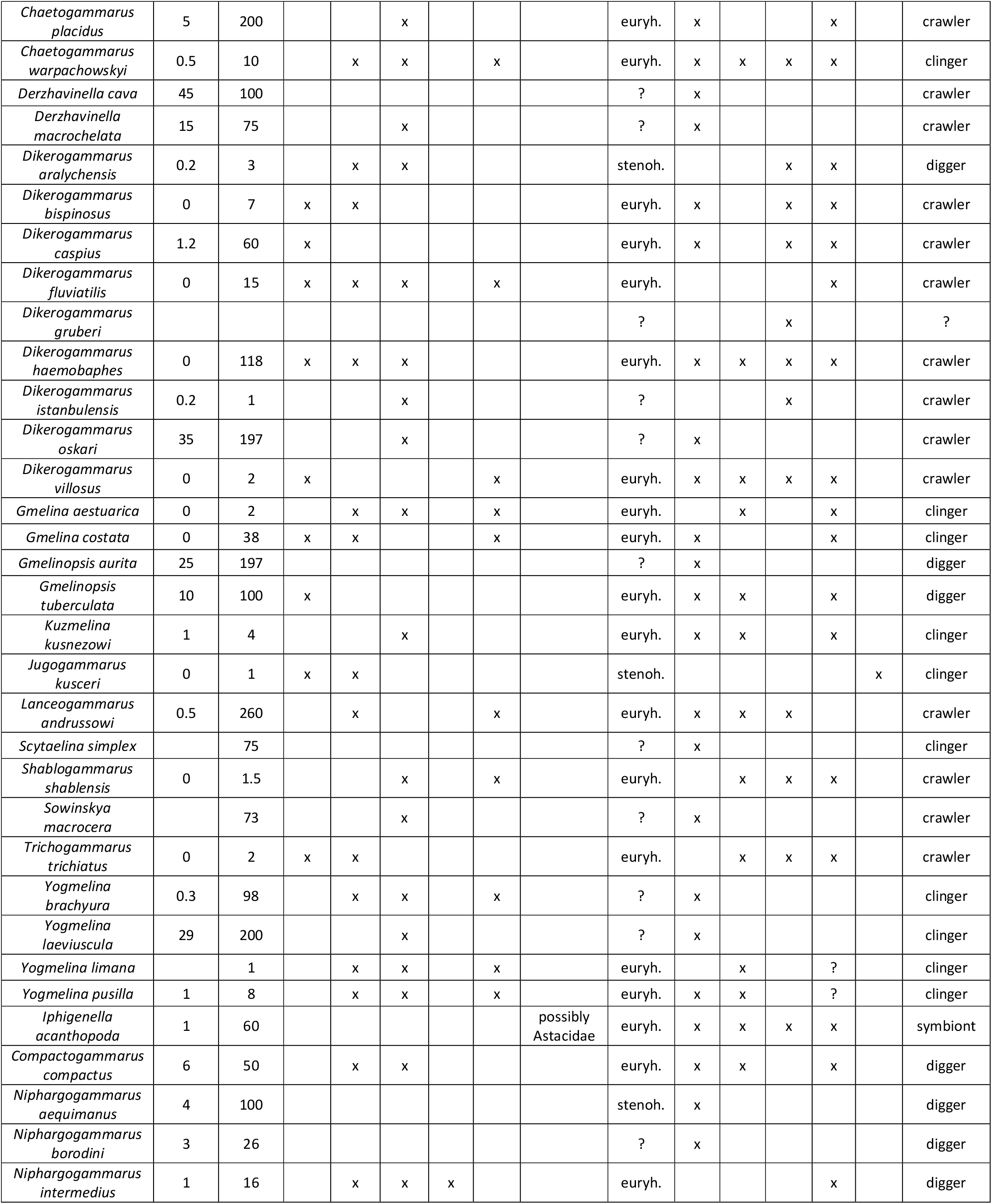
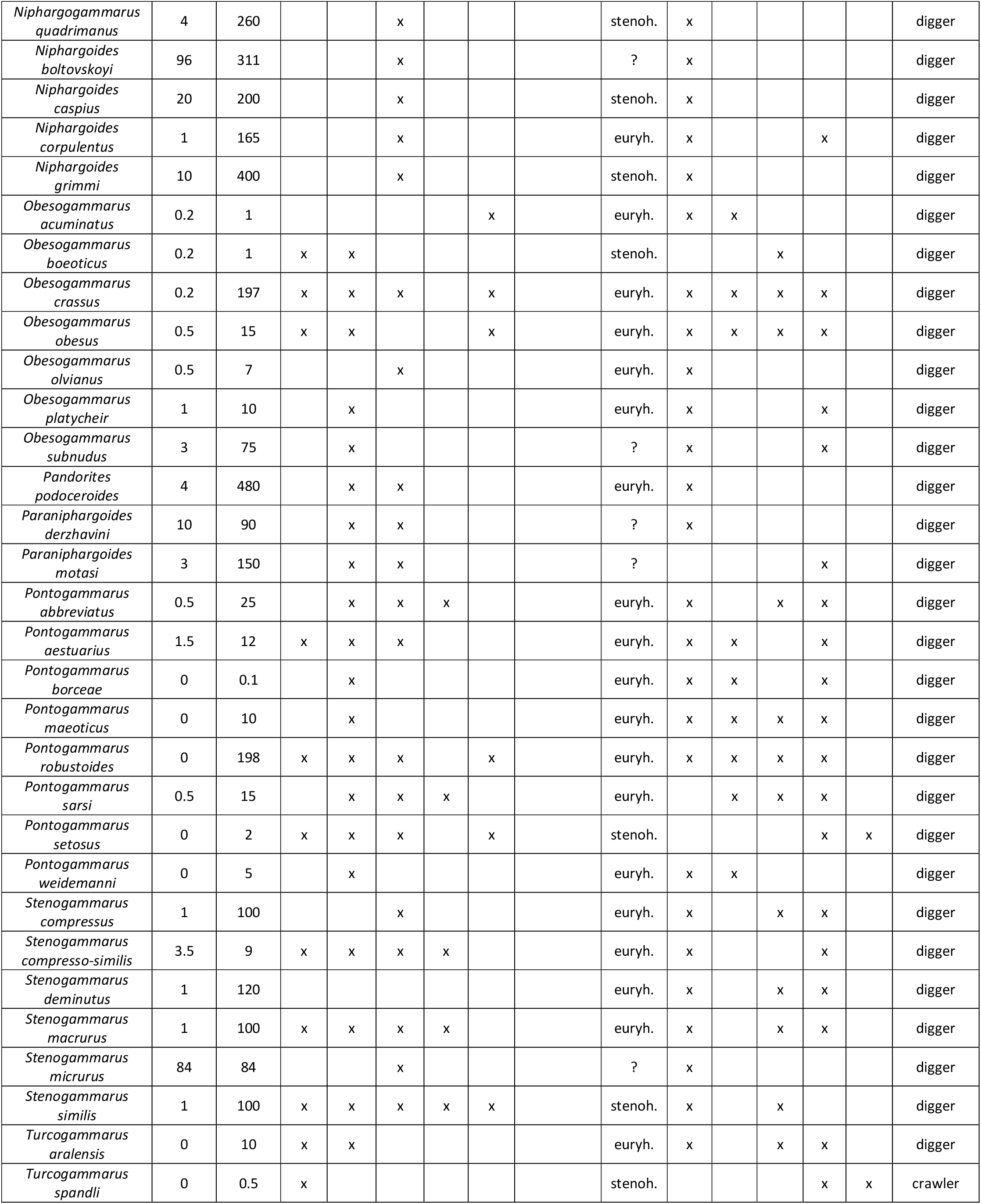
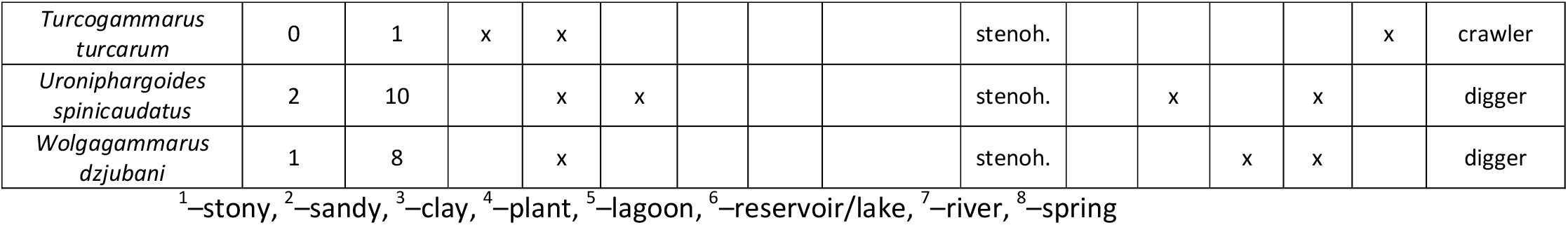
Ecological diversity of Ponto-Caspian gammaroid amphipods.

The depth gradient is broad, ranging from the wet sand of the supra-littoral to more than 500 m depth (Table 2, Fig. 5). Individual species also seem to be quite plastic and can be found from shallow depths (less than 50 m) to more than 200 m. The genera *Amathillina, Chaetogammarus, Niphargoides* and *Pandorites* have the broadest depth ranges. Species diversity is the highest in the first 50 m (79 species), then rapidly decreases to below 10 species in the 250-550 m interval (Fig. 5). The only species known to occur at depths greater than 500 m is *Chaetogammarus pauxillus*.

### Proposed ecomorphs

By integrating morphology and substrate type we aimed to classify the species into putative ecomorphs. Specifically, we looked for common morphological characteristics among taxa, while taking into account their similarity in PCA morphospace. We also took into account previous informal groupings of genera (Barnard and Barnard 1983). Once these groups were identified, their substrate preference was established by estimating the proportion of species occurring on a particular substrate. The substrate classification was simplified and divided into four groups: coarse (corresponding to stones and gravel), fine (corresponding to sand, mud and clay), plant and symbiotic. We acknowledge that this is a somewhat arbitrary approach. However, more sophisticated analyses could not be performed given the scarce data at hand. Quantitative data regarding ecology (substrate or trophic niche) are only limited to a few invasive species. Likewise, morphology is incompletely known in many species (especially mouthparts). We emphasize that our goal here was to provide a first exploratory step into understanding the connection between morphology and ecology.

We tentatively defined four ecomorphs: clingers, crawlers, diggers and symbionts. Loosely, these ecomorphs correspond with the currently recognized families and informal groupings of Barnard & Barnard (1983): crawlers with Gammaridae or “Echinogammarids” + “Dikerogammarids” (*sensu* Barnard & Barnard, 1983), clingers with Gammaridae or “Gmelinids” (*sensu* Barnard & Barnard, 1983), diggers with Pontogammaridae or “Pontogammarids” + “Compactogammarids” (*sensu* Barnard & Barnard, 1983), and symbionts with Behningiellidae, Caspicolidae and Iphigenellidae or “Cardiophilids” (*sensu* Barnard & Barnard, 1983). Below we describe the morphological and ecological peculiarities of each ecomorph.

1. Clinger. Stout body often keeled and/or ornamented with spines and tubercles, antennae are slender, short to medium length, coxal plates medium to deep, gnathopods weak, and pereopods short to medium length with pairs 3-4 strongly opposable to pairs 5-7 (Fig. 6). Clingers are intermediate in morphospace between crawlers and diggers, although there is significant overlap significantly with the latter group (Fig. 3b). Most species are associated with plants and fine substrate (Fig. 3b). Around 19% of all species belong to this ecomorph. Taxonomic composition is given in Table 2. Representative genera: *Axelboeckia* and *Gmelina*.
2. Crawler. Body is slender and generally smooth, antennae are long and slender, coxal plates shallow, pereopods slender, short to medium, gnathopods generally strong, and uropods long (Fig. 6). It is generally well-defined in morphospace having little overlap with clingers and diggers (Fig. 3b). Species are mainly associated with fine and coarse substrates (Fig. 3b). Around 26% of species belong to this ecomorph. Taxonomic composition is given in Table 2. Representative genera: *Chaetogammarus* and *Dikerogammarus*.
3. Digger. Stout body and almost exclusively smooth, antennae very short and thick, with 1^st^ article of antenna 1 often swollen, coxal plates deep, gnathopods generally strong, pereopods medium to long, with broadened articles often fringed with long and dense setae (Fig. 6). Diggers are very distinct in morphospace from crawlers and symbionts, but overlap noticeably with clingers (Fig. 3b). Species of this ecomorph predominantly occur on fine substrates and are characterized by a fossorial behavior (Fig. 3b). This appears to be the most common ecomorph since almost half of the Ponto-Caspian species are classified as diggers (49%). Taxonomic composition is given in Table 2. Representative genera: *Pontogammarus* and *Niphargoides*.
4. Symbiont. Very stout and generally minute bodies, with well-developed coxal plates and pereopod bases, usually characterized by diminished mouthparts (palps of maxilla 2 and maxilliped), pleon, urosome, antennae and pereopods (Fig. 6). The gnathopods can be very specialized (*Caspicola* and *Iphigenella*), or rudimentary (*Behningiella* and *Cardiophilus*). This ecomorph is the most distinct in morphospace, with hardly any overlap (Fig. 3b). Its species are known to live on or inside bivalve mollusks (*Cardiophilus* and *Caspicola*), or commensals with crayfish (*Iphigenella*). This ecomorph is the rarest and accounts for 6 % of all species. Taxonomic composition is given in Table 2. Representative genera: *Cardiophilus* and *Iphigenella*.

## Discussion

Our study reviewed and quantified for the first time the rich taxonomic, ecological and morphological diversity of Ponto-Caspian amphipods. Although we consider these findings preliminary, our synopsis will serve as a foundation for future eco-evolutionary and systematic studies. Below we discuss the evidence accrued so far that point towards a remarkable, yet unrecognized adaptive radiation. Within each of the following sub-sections we also highlight the gaps in existing knowledge and recommend further research.

### Ponto-Caspian gammarid amphipods – an adaptive radiation

The main prerequisites that define an adaptive radiation are: monophyly, species sympatry, speciation rate increase, and ecomorphological divergence (Schluter 2000; Simões et al. 2016). With respect to Ponto-Caspian amphipods the sympatry criterion is the most readily fulfilled since most of the species co-occur in the Caspian Sea and Lower Volga (Table 1). Furthermore, most species seem to be widespread in the Caspian Sea, occurring in all of its main areas (north, middle and southern) (Pjatakova and Tarasov 1996). A significant number of species are also found in sympatry in the Ponto-Azov region (Cărăuşu et al. 1955).

The monophyly condition is supported by recent molecular phylogenies which indicate that several morphologically disparate Ponto-Caspian genera form a well-supported clade (Copilaş-Ciocianu, Borko, et al. 2020; Hou et al. 2014; Hou and Sket 2016; Sket and Hou 2018). Although relatively few taxa have been sequenced so far, it is likely that the remaining species would fall within the same clade. The Ponto-Caspian amphipod radiation also satisfies the requirement of speciation rate increase since it experienced a higher diversification rate in comparison to its sister clades (Hou et al. 2014).

We consider that our current study fulfills, at least partially, the criterion of ecomorphological divergence, which is perhaps the most relevant to the adaptive radiation model. We highlight significant ecological and morphological disparity within the Ponto-Caspian amphipod radiation. Along an order of magnitude body-size gradient, morphology ranges from minute (several millimeters), stout-bodied symbiotic species with attenuated appendages, to large and slender (several centimeters), stocky and setose, or heavily armored species. Likewise, ecological diversity is also remarkable, with species being encountered along a >500 m depth gradient on virtually all types of substrates and water bodies (mountain springs to deep sea). By integrating morphology and ecology, we propose a provisional classification into four main ecomorphs: clingers, crawlers, diggers and symbionts. Although this classification is only tentative, we consider it a necessary first step towards understanding the evolution of Ponto-Caspian amphipods. We highlight that these ecomorphs have potential analogues in distantly related marine or Lake Baikal taxa that occupy similar habitats (see Morphological evolution section below), further strengthening the environment-phenotype association.

Overall, it appears that Ponto-Caspian amphipods fulfill, at least to some extent, the main prerequisites of the adaptive radiation model. However, our findings provide only a first glimpse. Extensive further research is needed to corroborate the patterns highlighted herein. Specifically, the criteria of monophyly and speciation rate increase have to be tested on larger multilocus phylogenies with a greater taxonomic coverage. The morphology-environment association needs to be refined with newly collected field data. Specifically, fine-scale morphometry of functionally relevant traits coupled with trophic niche (gut content DNA metabarcoding and stable isotopes) and ecology (depth, substrate and salinity) in a phylogenetic context will provide a more comprehensive ecomorphological understanding. Furthermore, it is important to test whether these ecomorphs have a common ancestor or evolved several times independently (Trontelj et al. 2012). It is likely that upon more detailed investigation they could be split into more specialized forms. Comparative transcriptomics and genomics could provide important insight into adaptation and selection at the molecular level. A well-sampled time-calibrated molecular phylogeny could also prove invaluable for understanding the historical circumstances that promote the evolution of invasive species.

### Morphological evolution

Recent molecular phylogenies revealed that the morphologically diverse Ponto-Caspian amphipod radiation is nested within the genus *Echinogammarus* (Hou and Sket 2016; Sket and Hou 2018), which is characterized by morphological conservatism (Pinkster 1993). This is in good agreement with previous hypotheses that postulated a close relationship between these two groups (Barnard and Barnard 1983). A similar pattern is also encountered in the two highly diverse Baikal amphipod radiations which are classified into several families (Hou and Sket 2016; Lowry and Myers 2013), yet they are both nested within the genus *Gammarus* (Hou et al. 2011, 2014; Macdonald et al. 2005; Naumenko et al. 2017), notorious for its low morphological diversity, morphological crypsis (Copilaş-Ciocianu and Petrusek 2015; Katouzian et al. 2016; Mamos et al. 2014) and generalist ecology (MacNeil et al. 1997; Piscart et al. 2011). And yet again the same pattern appears in the distantly related American genus *Hyalella* where morphologically conserved riverine species (Witt et al. 2006) colonized the ancient Titicaca Lake multiple times, giving rise to a remarkable array of forms (Adamowicz et al. 2018; González and Coleman 2002; Jurado-Rivera et al. 2020). These compelling patterns indicate that species living in ephemeral, highly fluctuating and ecologically limited environments (springs, streams, rivers and shallow lakes/ponds) are under stabilizing selection for maintaining a generalist life-style and a conserved, non-specialized morphology (Wellborn and Broughton 2008). On the other hand, species inhabiting stable ancient lakes with broad niche space are probably under disruptive selective pressures which in turn promote specialization and ecological speciation (Seehausen 2015; Wellborn and Langerhans 2015). Thus, it would seem that the ecological transition from ephemeral habitats to long-lived ancient lakes promotes adaptive radiations in some freshwater amphipod groups. These intriguing patterns are worth pursuing further and could shed more light on the role of ecological opportunity in driving adaptive radiations.

We propose that the ecological and morphological diversity of Ponto-Caspian gammarids can be distilled into four ecomorphs. Remarkably, all of them apparently have analogues in distantly related lineages inhabiting oceanic waters or other ancient lakes (Figs. 7-8). The Ponto-Caspian symbiotic ecomorph is the most specialized and morphologically distinct due to its reduced mouthparts, antennae, pereopods and urosome, presumably due to a semi-parasitic life-style. We highlight a striking resemblance between the Ponto-Caspian genus *Behningiella* and the oceanic algae-boring genus *Bircenna* Chilton, 1884 (Fig. 7a). Both exhibit typical features for substrate boring such as a large head with protruding mandibles adapted to cutting into tough material, and extremely short antennae and pereopods due to living in narrow self-constructed tunnels (Mejaes et al. 2015). Within the Baikal Lake Acanthogammaride radiation, the symbiotic ecomorph is probably represented by the parasitic genus *Pachyschesis* (Naumenko et al. 2017; Takhteev 2019).

**Fig. 7.**
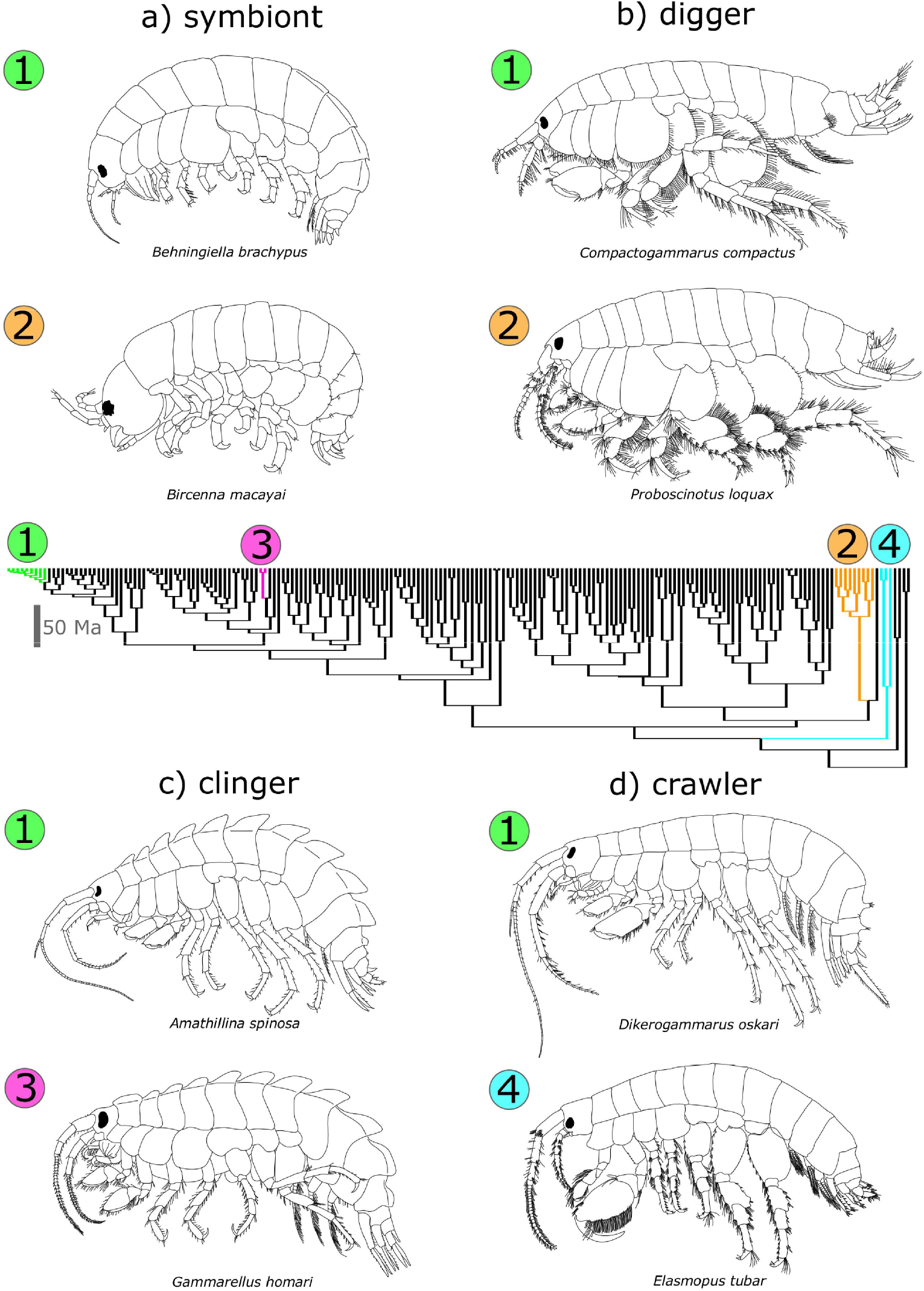
Putative examples of ecomorphological convergence of Ponto-Caspian and distantly related oceanic taxa. Ponto-Caspian species are shown with green. a) Symbiotic ecomorph adapted to piercing various organic substrates (redrawn from Derzhavin (1948) and Loerz et al. (2010)), b) digger ecomorph adapted for digging and burrowing in fine substrates (redrawn from Sars (1895) and Barnard (1967)), c) clinger ecomorph adapted to cling on algal and vegetal substrates (redrawn from Sars (1896)), and d) crawler ecomorph adapted to a generalist life-style, usually hiding in coarse stony substrates (redrawn from (Sars (1896) and Garcia-Madrigal (2010)). The phylogenetic tree is a time-calibrated molecular phylogeny of Amphipoda modified after Copilaş-Ciocianu et al. (2020)

**Fig. 8.**
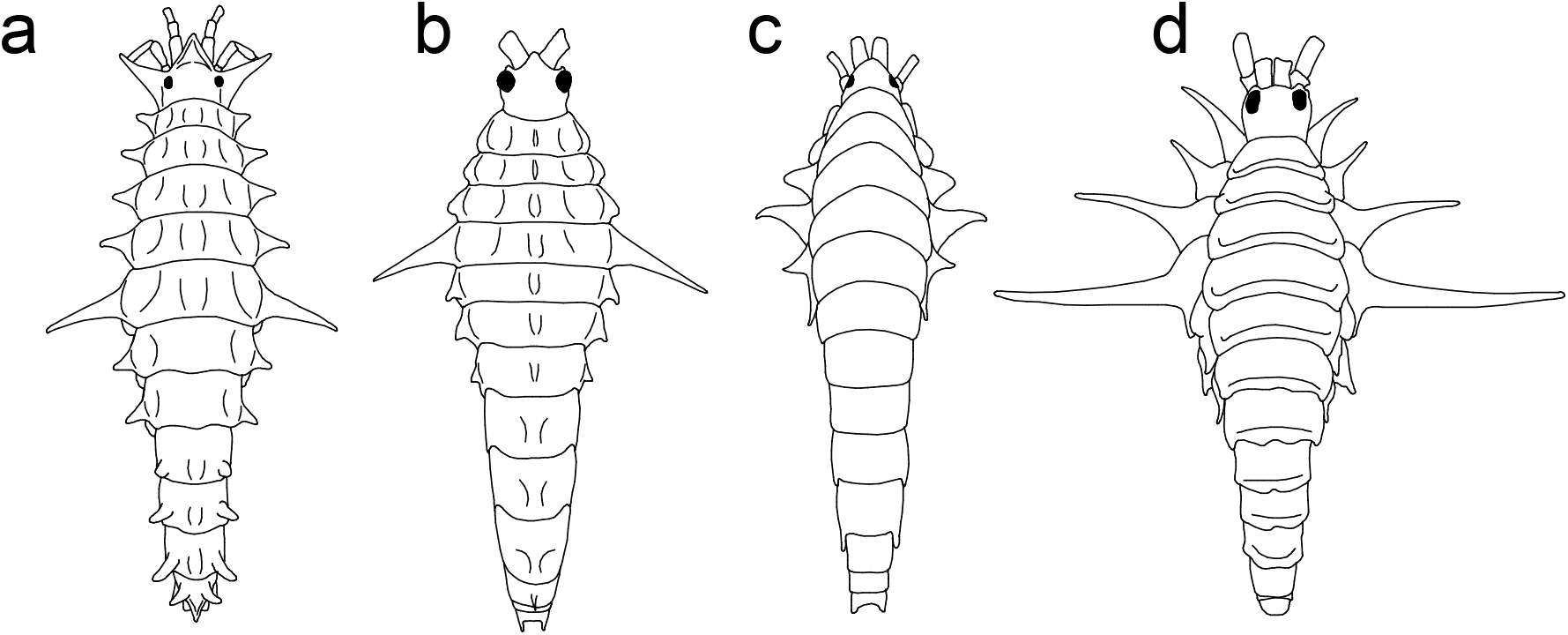
Examples of evolutionary convergent patterns in body armature of species inhabiting different ancient lakes. a) *Axelboeckia spinosa* (Caspian Sea, redrawn after Sars (1894b)), b) *Acanthogammarus lappaceus* (Lake Baikal, redrawn after Daneliya et al. (2011)), c) *Issykogammarus hamatus* (Lake Issyk-Kul, redrawn after Chevreux (1908)), and d) *Hyalella armata* (Lake Titicaca, redrawn after González & Coleman (2002))

The fossorial ecomorph seems to be the most common among Ponto-Caspian amphipods. These species are generally adapted for digging in fine substrates and have stout, strong bodies with very short yet powerful and thick antennae, and broadened pereopods usually fringed with dense rows of setae. This ecomorph is widely encountered among amphipods in general, albeit under slightly different iterations (Bousfield 1970). Morphologically, most fossorial amphipods are classified within the superfamily Haustorioidea (Lowry and Myers 2017). However, molecular phylogenies indicate that the fossorial body-type evolved multiple times independently (Copilaş-Ciocianu, Borko, et al. 2020; Hancock et al. 2020). A noticeable resemblance can be observed between the Ponto-Caspian genus *Compactogammarus* and the hyaloidean *Proboscinotus* Barnard, 1967 (Fig. 7b). Additionally, in Lake Baikal this ecomorph is possibly represented by the Micruropodidae radiation, comprising fossorial species living on fine substrate (Naumenko et al. 2017; Takhteev 2019).

The clinger ecomorph characterizes species with elaborate body armature/ornamentation and preference for living (plant) substrate. These species often have elongated and curved dactyls for improved grasping of the substrate. Given the exposed nature of their life-style, the armature might serve as protection against predators (Bollache et al. 2006; Copilaş-Ciocianu, Borza, et al. 2020) or, in combination with variegated coloration (as is often the case with armored taxa), may act as camouflage by disrupting the body contour (d’Udekem d’Acoz and Verheye 2017). We point out the high similarity among the Ponto-Caspian genus *Amathillina* and the oceanic algae-clinging *Gammarellus* Herbst, 1793 (Fig. 7c). Although the Ponto-Caspian clingers are diverse in ornamentation and armature, some striking resemblance can be observed with Baikal Lake taxa. For example *Amathillina* and *Eucarinogammarus* (Baikal), *Axelboeckia* and *Acanthogammarus* (Baikal), and *Kuzmelina* and *Propachygammarus* (Baikal) (Naumenko et al. 2017; Takhteev 2019).

The crawler ecomorph is the second-most encountered in Ponto-Caspian amphipods, characterizing species living on coarse or fine substrate, often in shallow water. Typically, these taxa are strongly sexually dimorphic, males possessing very large second gnathopods, relatively long antennae and slender bodies with shallow coxal plates. Morphologically, this morph is probably the most plesiomorphic, being widespread among the amphipod evolutionary tree, especially in some basal branches (Copilaş-Ciocianu, Borko, et al. 2020; Lowry and Myers 2017) as well as in the oldest known fossils (Jarzembowski et al. 2020). As an example, we emphasize the similarity among the Ponto-Caspian genus *Dikerogammarus* and the widespread littoral genus *Elasmopus* Costa, 1853 (Fig. 7d).The Baikalian analogues of this ecomorph could be envisioned in *Eulimnogammarus* and *Corophiomorphus* (Naumenko et al. 2017; Takhteev 2019).

Body armature is extremely diverse in amphipods, with similar phenotypes having evolved independently multiple times (Copilaş-Ciocianu, Borko, et al. 2020; Lowry and Myers 2017; Naumenko et al. 2017). We highlight a remarkably convergent evolution of body armature in some ancient lake radiations where strong lateral spines appear on the pereonites, the longest one being located on the 4^th^ or 5^th^ segment (Fig. 8). In some cases the spine is an outgrowth of the tergum, while in others an outgrowth of the coxal plate. These analogous convergent structures point towards a strong selective pressure. Most likely these spines function as a mechanism for deterring ingestion by predatory fish (Bollache et al. 2006; Copilaş-Ciocianu, Borza, et al. 2020), although the exact mechanical interactions are unknown.

### Spatio-temporal origin

The phylogenetic position of Ponto-Caspian amphipods within the Atlanto-Mediterranean *Echinogammarus* clade (*sensu* Hou et al., 2014; Sket & Hou, 2018) indicates that this radiation likely has a Mediterranean origin. Specifically, its sister clade is represented by the genus *Dinarogammarus*, which is endemic to freshwaters of the Western Balkans (Sket and Hou 2018). Regarding the temporal time-frame, several recent studies proposed a Middle Miocene origin (ca.12-14 Ma) (Copilaş-Ciocianu, Borko, et al. 2020; Hou and Sket 2016), coeval with the final closure of the Paratethys, which caused a switch from marine to brackish conditions and promoted the evolution of endemic faunas after initial mass extinctions (Popov et al. 2004; Rögl 1999). This time frame is also supported by Late Miocene (ca. 9-10 Ma) Caucasian fossil taxa (two genera and five species) that have clear affinities with extant Ponto-Caspian genera *Axelboeckia, Gmelina, Kuzmelina* and *Yogmelina* (Derzhavin 1927, 1941). Alternatively, an earlier study suggested an origin dating back to the Eocene (30-40 Ma) (Hou et al. 2014). However, this analysis was based on biogeographical calibration of the molecular clock rather than fossils, thus possibly resulting in biased inferences (Ho et al. 2015). Furthermore, a Late Eocene origin does not correspond with an isolation of the Paratethys realm from the world ocean (Popov et al. 2004). As such, we consider that a middle Miocene origin is more plausible considering the data at hand.

A densely sampled, multilocus and time-calibrated phylogeny will be of critical importance in understanding the historical biogeography and evolution of Ponto-Caspian gammarids. Furthermore, such a phylogeny could complement geological studies regarding the palaeogeographic history of the Paratethyan region, as seen with other freshwater gammarids (Copilaş-Ciocianu et al. 2019; Copilaş-Ciocianu and Petrusek 2017; Hou et al. 2011; Mamos et al. 2016). It could provide additional time constraints on some important palaeogeographic events such as the final Paratethys closure, the isolation of the Pannonian, Pontic and Caspian basins, the emergence of the Caucasus, as well as the recurrent episodic connections of the Pontic and Caspian basins during the Plio-Pleistocene.

### Taxonomic and systematic remarks

The Ponto-Caspian gammaroid amphipods as defined in this study are formally split into 5 families: Behningiellidae, Caspicolidae, Gammaridae, Iphigenellidae and Pontogammaridae. However, molecular research has revealed that Pontogammaridae is nested within Gammaridae, and also harbors the gammarid genus *Dikerogammarus* (Copilaş-Ciocianu, Borko, et al. 2020; Hou et al. 2014). Members of this family correspond to the digger ecomorph, which probably evolved on more than one occasion. Moreover, the Ponto-Caspian “Gammaridae” form a paraphyletic grade at the base of Pontogammaridae (Sket and Hou 2018). A taxonomically more inclusive morphological and molecular study will clarify this issue, but most likely will not recover Pontogammaridae as monophyletic. However, for the sake of stability we do not propose any taxonomic changes until this issue is firmly resolved.

The remaining families Behningiellidae, Caspicolidae and Iphigenellidae are poorly known and have not yet been sequenced. Behningiellidae and Iphigenellidae have been classified into Gammaroidea based on a morphological cladistic analysis (Lowry and Myers 2013). However, the monotypic Caspicolidae is currently not recognized as part of Gammaroidea, but as a distinct superfamily (Caspicoloidea) within the infraorder Talitrida (Lowry and Myers 2013). This classification is erroneous because the authors mistakenly considered that the antenna I lacks an accessory flagellum (a defining character state of the infraorder Talitrida). Derzhavin’s (1944) original description clearly indicates the presence of the accessory flagellum, although it is reduced and uniarticulate. Another issue with assigning Caspicolidae to Talitrida is the presence of a well-developed mandibular palp, whereas an absent/vestigial palp is another defining character state of the Talitrida (Lowry and Myers 2013). Behningiellidae, Caspicolidae and Iphigenellidae belong to the symbiotic ecomorph and represent highly specialized taxa which are difficult to classify using external morphology alone. It is very likely that these small families are nothing but highly derived Ponto-Caspian gammarids, possibly related to the various genera of the gmelinid facies (*Gmelina, Kuzmelina* and *Yogmelina*) (Barnard and Barnard 1983; Bousfield 1977; Derzhavin 1944). Thus, the systematic position of these families will be clarified only with additional morphological and molecular study.

We argue that most, if not all Ponto-Caspian amphipod species are in need of thorough, modern revision using morphology, multilocus DNA sequences and ecology. Many species are only partially illustrated and intraspecific variability has been studied in only a handful of taxa (Cărăuşu 1936; Nahavandi et al. 2013). Moreover, cryptic lineages of potential specific status have been recently discovered (Jażdżewska et al. 2020). As such, a first step towards a modern taxonomic revision could be the generation of a well-sampled DNA barcode reference library.

Lastly, the Late-Miocene (Upper Sarmatian, ca. 9 Ma) fossil genera *Andrussovia* and *Praegmelina* have long been considered ancestral to extant Ponto-Caspian genera such as *Gmelina* and *Amathillina*, albeit without a formal analysis (Barnard and Barnard 1983; Derzhavin 1927, 1941). The fossils were discovered in calcareous clay deposits at the foothills of the Caucasus near Grozny, Solenaya balka (Chechnya, Russian Federation) and from Eldar Oyugu Ridge (Azerbaijan). We agree that there are rather clear affinities with extant Ponto-Caspian species in general, mainly in the combination of the following traits: shape of the basis of pereopod 7, ornamentation and armature, short and thick antennae, and deep coxal plates. Some traits are considered plesiomorphic, such as the lack of a postero-ventral lobe on the basis of pereopod 7, and the long endopod of uropod 3. A cladistic analysis is necessary to confidently assess evolutionary relationships with extant taxa. Until then, these species should be conservatively treated as stem Ponto-Caspian amphipods (Copilaş-Ciocianu, Borko, et al. 2020).

Two more Miocene fossil taxa have been reported from the Caucasus that have less clear affinity with extant Ponto-Caspian taxa. These are *Gammarus praecyrius*, Derzhavin, 1941 and *Hellenis saltatorius*, Petunnikoff, 1914. The former is indistinguishable from a typical *Gammarus* and it is thus not considered a Ponto-Caspian taxon. The affinities of the latter taxon are less straightforward to interpret due to its high degree of morphological specialization (very short antennae, large raptorial gnathopods and unusually long pereopods). Such a combination of traits is not present in the extant Ponto-Caspian fauna. Furthermore, Petunnikoff’s illustrations are also not detailed enough to draw a conclusion. At the moment we consider that it is possible that *H. saltatorius* could be related to Ponto-Caspian amphipods but further detailed studies are needed.

## Conclusion

The Ponto-Caspian gammarid radiation fulfills, at least partially, the most important criteria of an adaptive radiation: 1) apparent monophyly, 2) sympatric occurrence within a constrained area, 3) accelerated diversification and 4) ecomorphological disparity. Nevertheless, these literature-based results are only preliminary and a lot of in depth eco-evolutionary study is further needed. Moreover, most species need a modern taxonomic revision within an evolutionary context. Nevertheless, we consider that Ponto-Caspian amphipods could be an excellent future model for the study of adaptive radiation, origin of invasive species, and could even help illuminate the region’s dynamic palaeogeographic history.

## Acknowledgements

This study was financed by the Lithuanian Research Council (contract no. 09.3.3-LMT-K-712-19-0149). We thank Nadezhda Berezina for providing important literature.

## Appendix

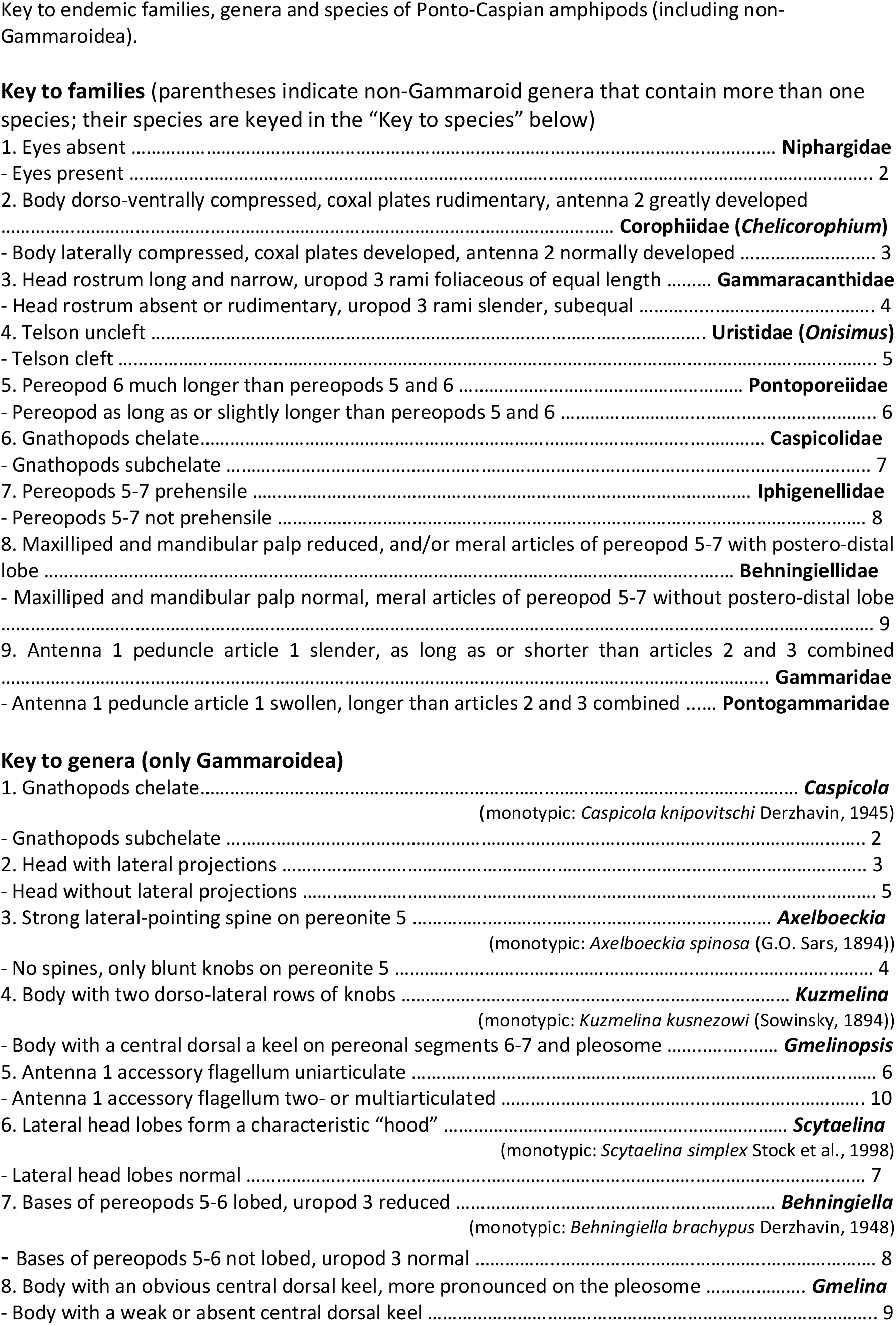
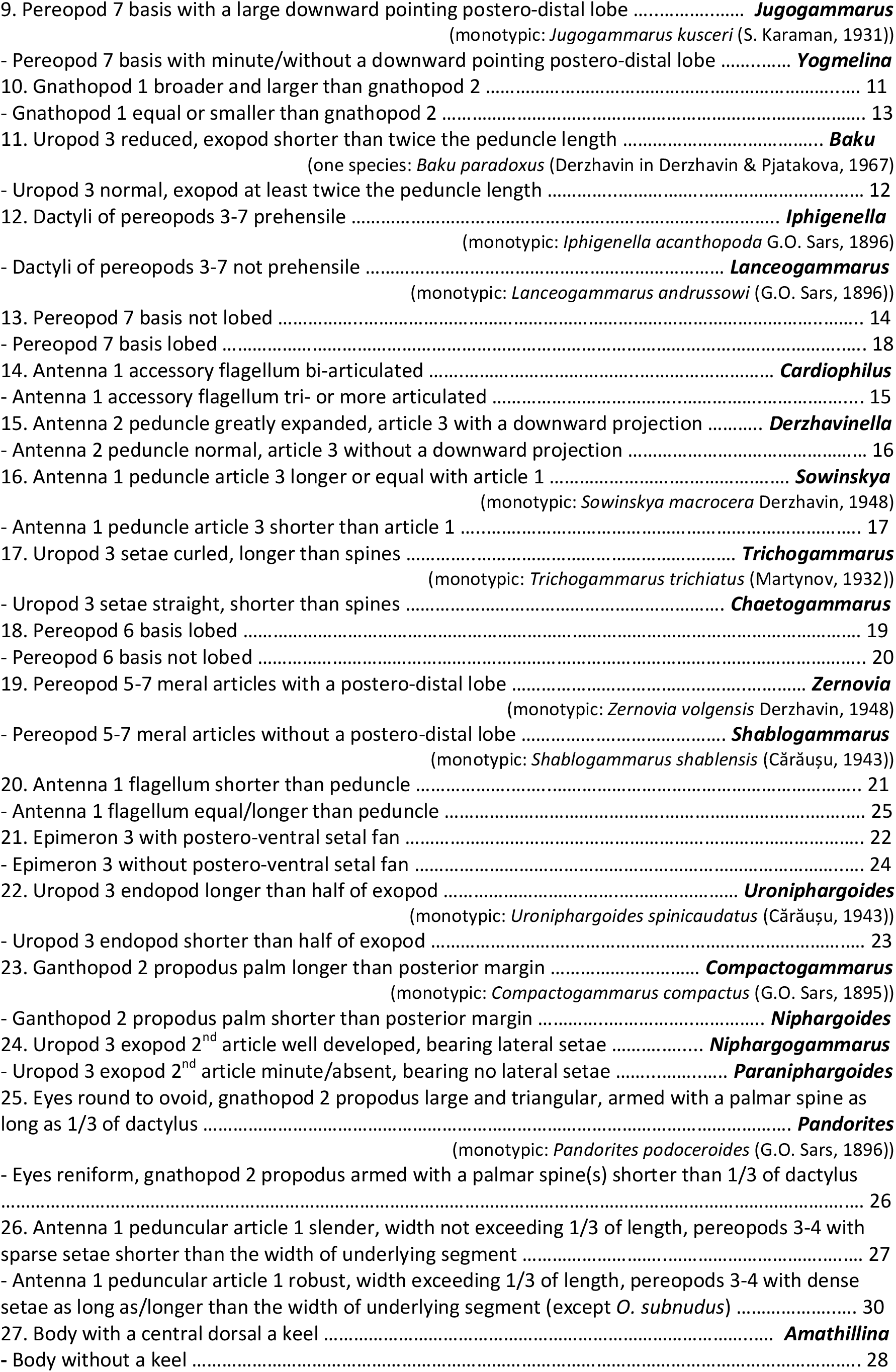
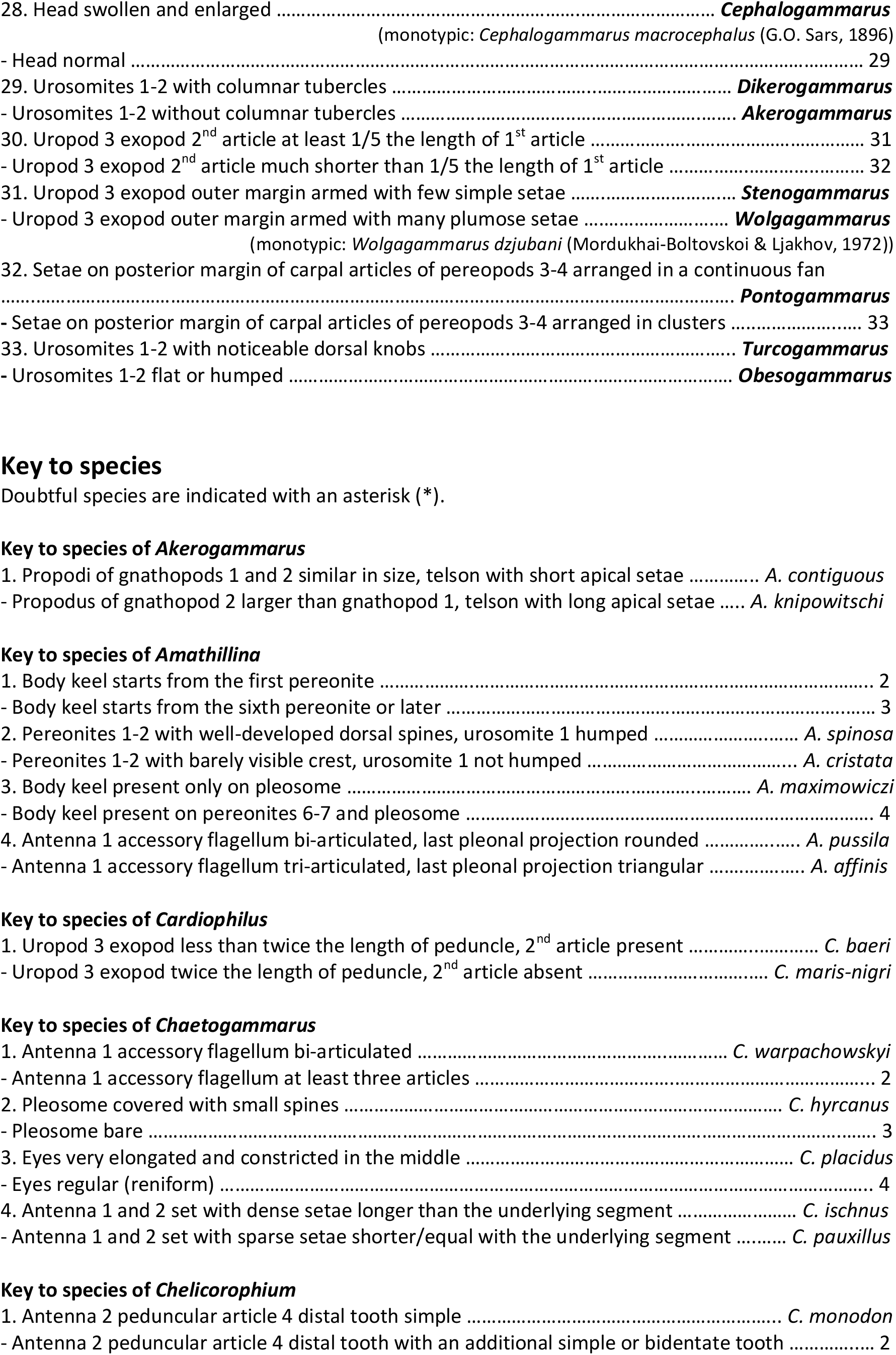
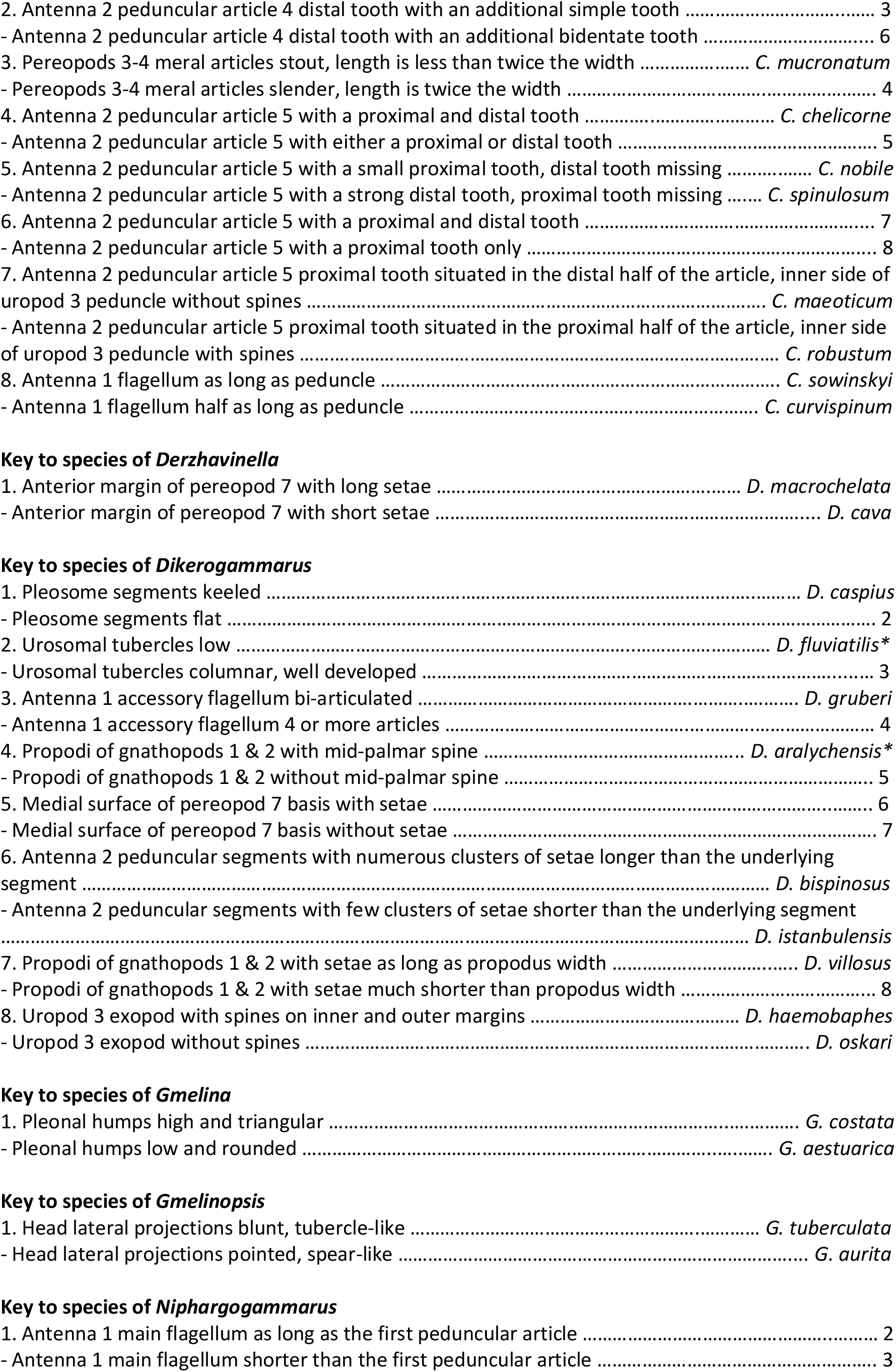
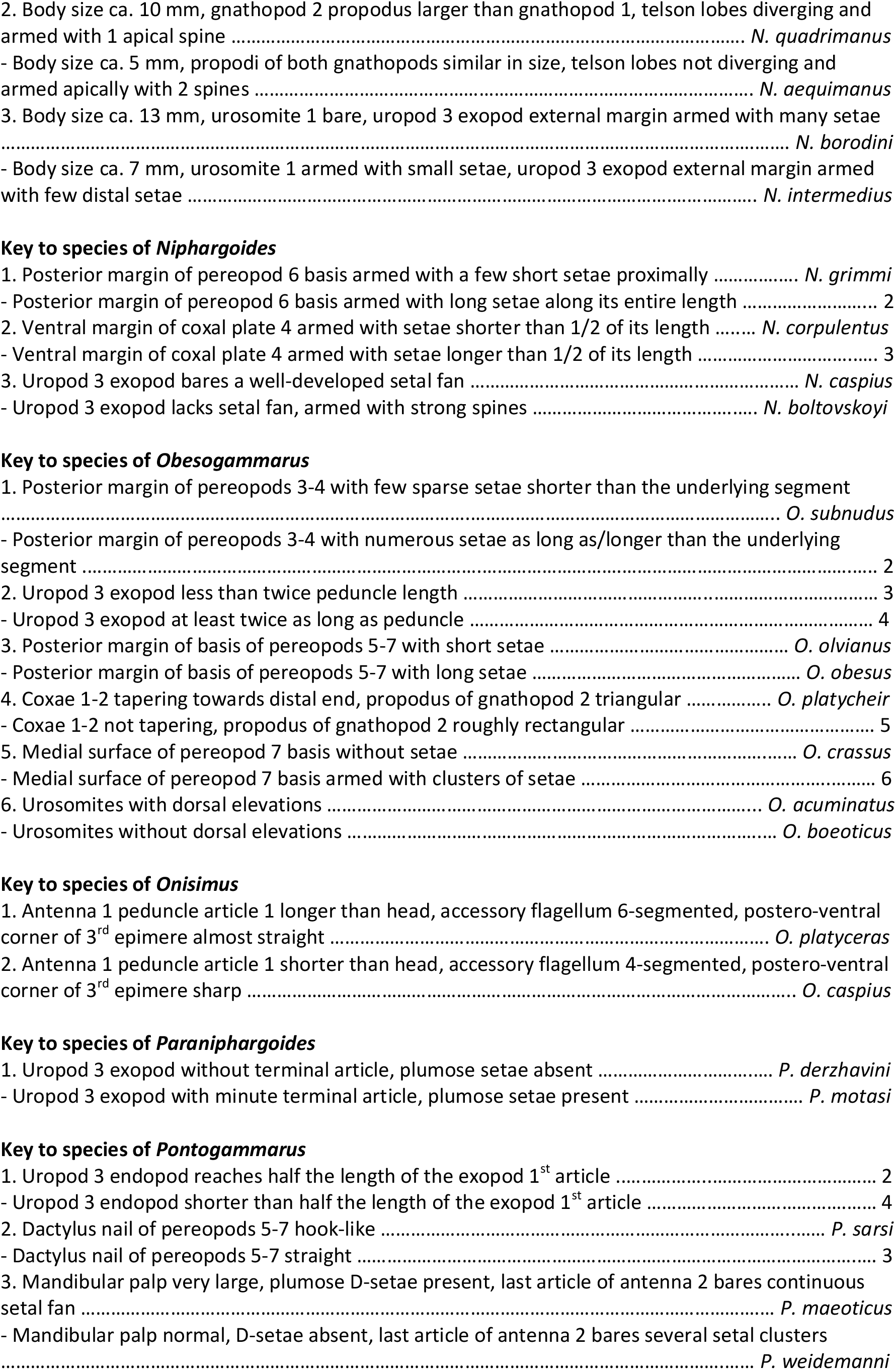
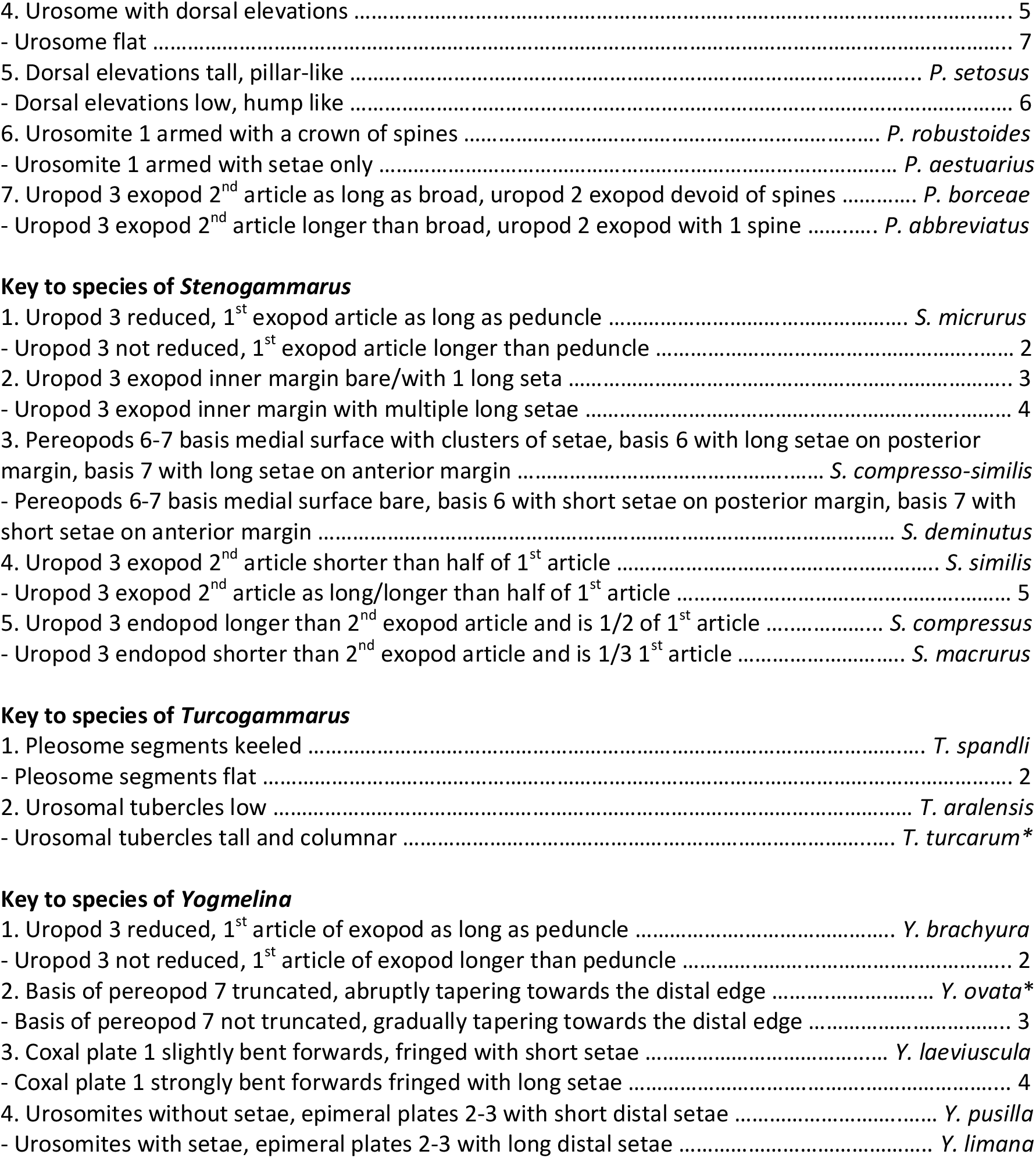

