## Supplementary figures and images for "Taxonomic, ecological and morphological diversity of Ponto-Caspian gammaridean amphipods: a review"

### Fig. S1

Numbers correspond with traits in Table S1

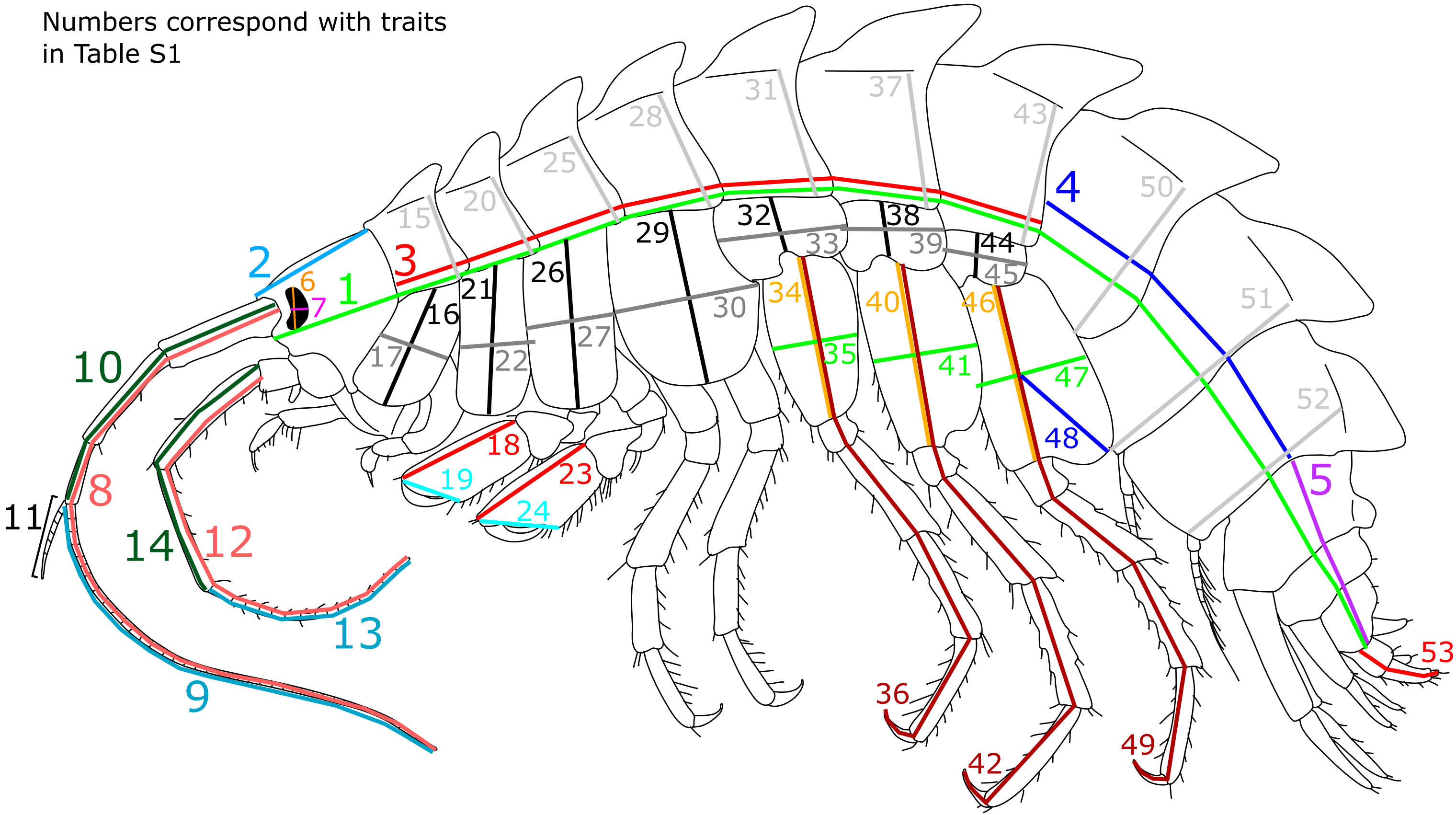
